# Transcriptional profiling of populations in the clam *Ruditapes decussatus* suggests genetically determined differentiation in gene-expression along parallel temperature gradients and between races of the Atlantic Ocean and West Mediterranean sea

**DOI:** 10.1101/2023.03.28.533615

**Authors:** Carlos Saavedra, Massimo Milan, Ricardo Leite, David Cordero, Tomaso Patarnello, Leonor Cancela, Luca Bargelloni

## Abstract

Ongoing ocean warming due to climate change poses new challenges for marine life and its exploitation. We have used transcriptomics to find genetically-based responses to increased temperature in natural populations of the marine clam *Ruditapes decussatus* living along parallel thermal gradients in southern Europe. Clams of the Atlantic and West Mediterranean races were collected in a northern (cool) and a southern (warm) localities. The animals were kept in running seawater in the warm, Atlantic southern locality for a 15-week period, during which water temperature raised to typical southern European summer values. After this period, the expression profile was obtained for a total of 34 clams and 11,025 probes by means of an oligonucleotide microarray. We found distinct transcriptional patterns for each population based on a total of 553 differentially expressed genes (DEG), indicating innate differences which probably have a genetic basis. Race and latitude contributed significantly to gene expression differences, with very different sets of DEG. A gene ontology analysis showed that races differed mainly at genes involved in ribosomal function and protein biosynthesis, while genes related to glutathione metabolism and ATP synthesis in the mitochondria were the most outstanding as to north/south transcriptional differences.

## 1. Introduction

The changes in the marine environment caused by the increase of CO_2_ in the atmosphere, mainly increased water temperature and reduced pH, have led to a growing interest in the physiological and molecular mechanisms that marine organisms can employ to face those changes [1–5]. In addition to traditional physiological and molecular biology approaches, genomic techniques are playing an important role in studying the interactions between organisms and the changing environment [1]. In particular, studies of gene expression at the whole genome level (transcriptomics) have been especially effective in showing the amount and types of genes, and the molecular networks, that are involved in the response to the change in seawater temperature and adaptation to it (e.g.: [6,7]). Determining which genomic regions and genes show genetically-based differential transcriptional responses is especially important in species of conservation or economic interest, because it can help to achieve a deep and focused research on the behavioural, physiological, cellular, biochemical and molecular mechanisms involved.

The use of transcriptomics in the framework of population biology has several complications. The first one is that the organisms are responsive to environmental changes by modulating the expression of their genes (plasticity). Therefore, the observed differences among individuals collected in different locations or time points can be due to environmental differences across their ranges, and also to temporal variation in the environment, such as seasonal variation. The second complication is that individuals from populations that are genetically differentiated can show differences in gene expression due to DNA variants in regulatory regions. Disentangling the effects of environment and genes on gene expression can be achieved by studying individuals from separate populations in one or more common environments, so that the relative effect of each source of transcriptomic variation can be established [8–12].

One of the clear results that have emerged from studies of population transcriptomics in relation to temperature is that individuals native to different localities often show differences in the patterns of gene expression when they are exposed to a common thermal environment in the laboratory or in the field [7,13,14]. This is expected, because organisms usually show genetic differences among populations. It has been shown by several studies that even slight changes in allelic frequencies at genetic markers may be associated to differences in expression patterns among populations [15]. These differences could affect, at least partly, their response to temperature changes. The differential responses observed may have a genetic basis, i.e.: they are caused by differences in the DNA sequence of the regulatory regions or master regulatory genes (e.g.: [16]).

One additional step in discovering the genetic basis of transcriptional responses to thermal change in the sea, is to know whether the populations living in contrasting thermal environments show specific adaptations to them. The observation of correlations of transcriptomic differences with differences in specific environmental factors of the native habitats is suggestive of adaptive responses (i.e.: responses based on genetic differences caused by natural selection). Latitudinal thermal gradients in the sea are nice natural experimental settings for testing adaptation, and several studies have used them to infer the adaptive nature of some gene expression patterns [17–21].

On the other hand, population genetic studies using genetic markers have shown that many marine species harbor cryptic geographic variability at the level of the DNA. This variability could result in different responses to environmental change across populations. In particular, studies of intraspecific phylogenies of DNA sequences (especially mitochondrial DNA) have shown the existence of particular phylogenetic lineages that are geographically restricted, separating groups of populations across regions that have evolved independently during long time periods, and are often separated through abrupt genetic change areas. These areas have been termed phylogeographic breaks [22]. The amount and characteristics of intraspecific physiological variation across phylogeographic breaks has been rarely studied in a systematic way [23], and gene expression variation has had the same destiny, with very rare exceptions [24]. We would expect transcriptomic differentiation to be greater across phylogeographic breaks than that observed between populations within each phylogeographic group, but we cannot tell because we simply have no data. Testing this hypothesis is of real interest, because it can help to better understand the process and the causes of genetic differentiation across phylogenetic breaks (e.g.: neutral vs. adaptive), and also can be useful for an adequate management of genetic resources in commercial or endangered species [9]. Specifically, the study of the transcriptomic differences related to thermal environments across phylogeographic breaks can shed light to the different ability of populations separated by the break to cope with seawater temperature increase due to climate change, and allow discovering genes and gene networks that are acting for this purpose but remain hidden to the researchers by the effect of phylogeographic differentiation.

In this paper we report on a study of gene expression differences among populations of the grooved carpet-shell clam (*Ruditapes decusatus*), a species that shows a subdivided population genetic structure as described above. The grooved carpet shell clam (GCS clam from now on) is an infaunal bivalve that lives in sandy bottoms of the Mediterranean and nearby Atlantic coasts (MEDAT), from Morocco to Norway. The species is highly appreciated as seafood, and aquaculture practice is now increasing in some countries [25]. However, most of the clam production still come from harvested natural beds, especially in southern Europe and northern Africa [26–29]. The range of the species extends along wide climatic gradients spanning the European and northern African coasts, which makes this species a nice subject to study thermal adaptation. On the other hand, genetic studies using genetic markers such as mitochondrial DNA sequences, intron restriction-fragment length polymorphisms (iRFLP) and microsatellites have shown that there is a marked genetic subdivision across the well-characterized Eastern-Western Mediterranean phylogeographic break [30]. Additionally, strong genetic differentiation exists between the also well characterized Atlantic -Western Mediterranean phylogeographic break, with some markers attaining F_ST_ distance values of over 0.3 [30,31]. As the planktonic larval phase of the GCS clam lasts for 8-15 days (Borsa and Diter, cited in [26] ; [32]), allowing for extensive gene flow, the observed amount of genetic differentiation among populations is unusual, and makes this species one of the most genetically diverse organisms so far described in the literature across the MEDAT [33,34]. As for species with similar patterns of genetic subdivision, the three genetic units (races, from now on) found in the GCS clam are thought to be the result of the species population subdivision during glacial periods in the Pleistocene [30,31].

In this study we have taken advantage of the parallel temperature gradients existing at both sides of the Iberian Peninsula to disentangle the effects of phylogeography and thermal environment on differences in gene expression among GCS clam populations in response to the normal temperature increase that occurs during spring in the waters of the Mediterranean Sea and the nearby Atlantic Ocean. Clams were sampled from four localities: a cool, northern and a warm, southern locality, in each marine basin. Then they were transported to one of the warm southern localities for acclimation, as sea warming is the expected outcome of climate change in a near future. They were exposed to nearly natural conditions, and experienced the natural gradual increase of temperature that occurs during spring in southern European waters before they were subjected to transcriptomic analysis by using an oligonucleotide microarray, with the aims of 1) quantifying the relative differences in gene expression between races and between populations living in different thermal regimes in response to natural temperature increase, and 2) discovering functional units (genes, gene networks) showing differential response to the same environment.

## 2. Materials and Methods

### 2.1. Clam sampling and experimental setting

Adult GCS clams of average size 30 mm were sampled from four localities in Spain, France and Portugal (Figure1). These localities were chosen as representative of the ATL and WMED genetic races of clams described by [30], and of warm (south) and cool (north) environments along the parallel thermal gradients of seawater typical of the Mediterranean and nearby Atlantic region (MEDAT). Additionally, 20-30 animals were collected for genotyping in each locality. Clams from AN, MN and MS were transported live to the IATS facilities upon collection, and individually labeled with a number code on the shell using a black marker. Clams were placed in tanks with running, sand-filtered seawater, and fed a mixture of microalgae (*Tetraselmis svecica, Isochrysis galbana*, and *Chaetoceros calcitrans*).

**Figure 1.**
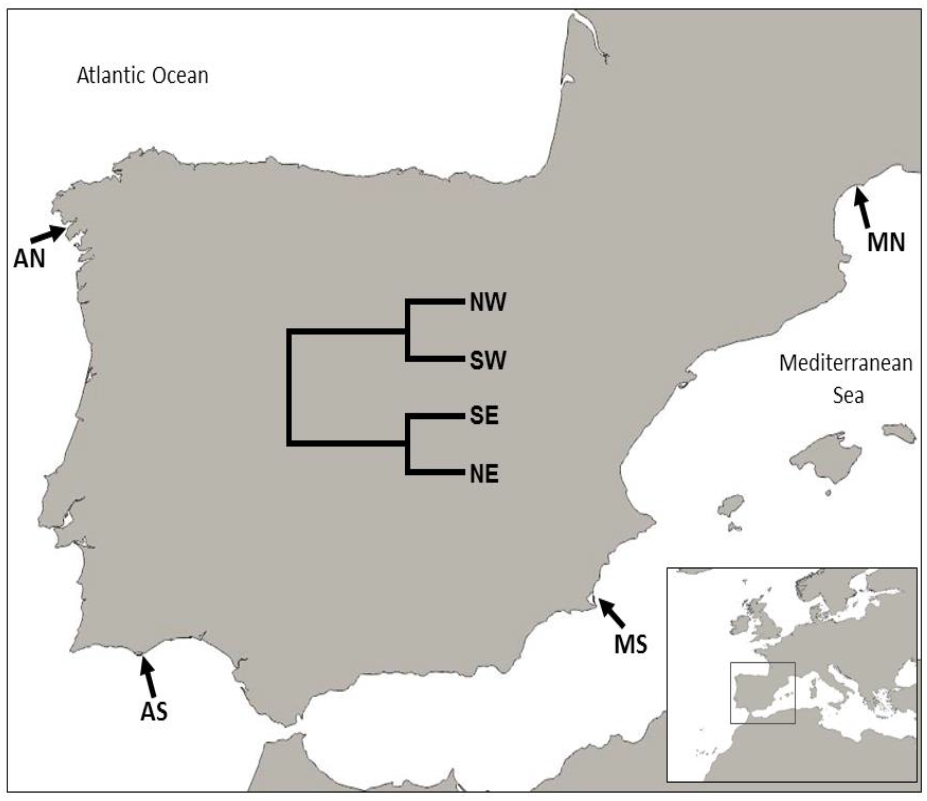
Map showing the localities sampled, and the NJ tree (inset) based on FST values computed from 6 intron-RFLP genetic markers. AN: Ría de Noia estuary (Spain). AS: Ria Formosa lagoon (Portugal); MN: Mar Menor lagoon (Spain). MN: Thau lagoon (France). A neighbor-joining tree based on six iRFLP is also shown.

On February 2012, 15 clams from AN, MN and MS were transported live in a cool container to the Ramalhete Marine Station at the University of Algarve near Faro (Portugal). Upon arrival clams were placed in an outdoor, 300 L, PVC tank provided with unfiltered running seawater. At this stage, 15 clams which had been collected a week before in the nearby shellfish beds of Ria Formosa, were incorporated to the experiment after being individually marked. They constitute the AS sample. The seawater was pumped directly in the tank from the estuary and therefore provided the clams with microalgae and suspended particles that constitute their usual diet in the wild. The tank was covered with a nylon mesh to avoid direct sunlight exposure and protect them from potential predators. Clams were inspected daily for mortality, and the condition of the setup was checked. Dead clams were recovered and discarded.

After a period of 14 weeks, all surviving clams were sacrificed by opening the valves with a scalpel. Gills were immediately dissected and preserved in Trizol at -80C. Mantle samples were preserved in ethanol 90%.

### 2.2. Environmental variables

Water temperature and salinity for each sampling locality were collected from a variety of sources. For the AN site, the weekly records for the Freixo station at -1 m for the period 2003-2009 were downloaded from the Instituto Tecnolóxico do Mar web site (http://www.intecmar.gal/Ctd/Default.aspx). Data for AS were obtained from the Estaçao Marinha de Ramalhete in Faro (Portugal). Data for MS come from the Oceanographic Information System of the Murcia region (https://siom-murcia.hub.arcgis.com/). These data were recorded at -1 m in an aquaculture site located at the same latitude as the sampling site but ca. 10 miles offshore, for years 2004-2010. Finally, the data for MN were obtained from REPHY-IFREMER (https://wwz.ifremer.fr/lerpc/Activites-et-Missions/Surveillance/REPHY). In this case, surface (0-1 m) data from the Bouzigues station were used. For AN, AS and MN, data were obtained once a week for a three-year period immediately before the experiment (2008-2010). In the case of MS data were available only as scattered records along the year, and the whole available series from 2004 to 2010 was used. The size of the sampled clams suggest that age could not be older than 3 years, so only data from years 2008-2010 were used for AN, AS and MN. During the course of the experiment, temperature and salinity at AS were measured as described.

### 2.3. DNA and RNA extraction

RNA was extracted at the University of Algarve laboratory with Trizol, following the protocols provided by the manufacturer. DNA from mantle samples preserved in ethanol were extracted with Chelex resin at the IATS-CSIC. RNA and DNA quality were checked by electrophoresis in agarose gels and spectrophotometric absorbance ratios. Concentrations were measured with a Nanodrop instrument. RNA integrity was examined with Bioanalyzer (Agilent).

### 2.4. Intron genotyping and population genetics

The polymorphism at six introns that allow the distinction of Atlantic and West Mediterranean races of GCS clam was scored by a simple technique based on PCR amplification of the introns followed by digestion with restriction enzymes and agarose gel electrophoresis [30].

iRFLP data were converted in genotypes. These data were pooled with those of 12 other populations previously studied [30]. Allele frequencies, heterozygosity and F_ST_ indices were calculated using GenePop [35]. A Bayesian analysis of genetic structure [36] was carried out to check that the samples collected showed the expected genetic profile of the ATL and WMED races. Details of this analysis can be found in [30]. A Neighbor-joining tree [37] was constructed from pair-wise F_ST_ among the four experimental populations.

### 2.5. Microarray hybridization

RNA from 41 individuals (15 AS, 9 MS, 12 AN and 5 MN) were hybridized to the one-color Agilent microarray previously designed from mRNA sequences obtained by RNA-seq from multiple organs and life stages of the species [37]. Eight slides were used, each containing 8 arrays. To minimize statistical confounding effects of slide, individuals from different populations were used to hybridize arrays in several slides, so that no slide had samples from less than 3 populations, and individuals from a population were scored together with individuals from at least 2 additional populations (due to sample size differences among populations, it was not possible to have all populations represented in all slides). Hybridizations were carried out in two batches of three slides. Six individual hybridizations that gave poor quality in the first hybridization batch were repeated in the second batch. At the end, two samples from AS, two from AN, and one from MN were discarded due to poor quality hybridizations, resulting in a total sample size of 36 clams. Normalized fluorescence data for these comparisons have been deposited in the GEO database under accession GSE223000.

### 2.6. Microarray data treatment and analysis

Fluorescence signal of reporter probes was checked for consistency with quality parameters following the recommendations of the microarray manufacturer. Fluorescence data were normalized by the Lowess method before analyses by using an in-house R script [39]. To ensure a minimum sample size in the analysis, and to avoid large variation in the sample size across probes that could lead to increased inter-probe variance, data were filtered by choosing only those probes that showed signal in at least 75% of the clams in each population. A first exploratory analysis of all filtered genes and samples using principal components was carried out to detect outliers. Fold change for each gene in each population was calculated as the ratio of the gene’s average population signal to the total sample average.

Normalized fluorescence intensity shown by each gene was first subjected to a one-way analysis of variance in order to test for the existence of significant expression differences among populations. The model for this ANOVA was

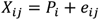

where *X*_*ij*_ stands for the fluorescence value of the *i* individual in the *j* population, *P*_*i*_ is the average fluorescence in population *j*, and *e*_*ij*_ is the error term.

Afterwards, a two-way analysis of variance was carried out to test for differential effects of fixed factors Race (Atlantic vs. Mediterranean) and Latitude (north vs. south) on individual gene expression. For each gene, the model for this ANOVA was

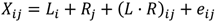

where *X*_*ij*_ stands for the fluorescence value of the i individual in the *j* population, *L*_*i*_ stands for the average effect of latitude *i, R*_*j*_ represents the average effect of the *j* race, (*L* · *R*)_*ij*_ represents the interaction between Race and Latitude and *e*_*ij*_ is the error term.

For all these analyses the TMeV software suite was employed [40]. P-values from ANOVA for each gene were used to estimate the false discovery rate by the Benjamini-Hochberg method [41]. Data were also subjected to hierarchical cluster analysis [42] in the R environment, using mean centered data, Euclidean distances, and the complete linkage option.

### 2.7. Functional analysis

Functional annotation of the sequences from which the probes were derived was carried out as described [38]. First, BLAST hits and Gene Ontology (GO) terms associated to probes were retrieved with Blast2GO [43], which makes a rigorous evaluation of the possible GO terms associated with the BLAST hits of each probe, and ascribes an index of certainty to each annotation. Valid annotations were considered if the index was higher than 0.5, using a threshold of 10-3 for BLASTx searches, and default parameters in the mapping and annotation steps. Overrepresentation of particular functional GO terms (i.e.: enrichment) in the groups of probes which were significant in the ANOVA sets of significant genes was examined by means of the Fisher exact tests using the DAVID platform [44] as described in [38], based on the GO terms associated with homologous genes of the zebrafish (Danio rerio). The use of this species is justified because it was the one to which BLAST analysis of the clam sequences produced the highest number of hits with annotations [38]. We did not intend to make overrepresentation tests a touchstone for the implication of differentially expressed genes and their associated GO terms in the functional aspects discussed in the text, and therefore we examined overrepresented terms with associated P-values slightly higher than 0.05 when they were supported by high values of other parameters such as fold-enrichment or count numbers.

### 2.8. Neutrality tests

Testing neutral or selective patterns of gene expression variation among populations was carried out for each gene using the “expression variance and evolution” model for quantitative trait evolution developed by Rohlfs and Nielsen [45], with the program EVE developed by the same authors.

## 3. Results

### 3.1. Environmental variables

Seawater temperature values recorded in the sampling sites in the three years previous to the study are summarized in Figure 2A. Average T values were lower in the northern populations (14.7 ± 0.3 ⁰C in AN, and 16.4 ± 0.5 ⁰C in MN), as compared to their southern counterparts in the same basin (18.9 ± 0.4 ⁰C in AS, and 19.5 ± 1.0 ⁰C in MS). Average values in Mediterranean sites were higher than in Atlantic sites. Maximum values were detected in August in all populations, and were over 27 ⁰C in all but the AN population, where the maximum recorded temperature was 19.9 ⁰C. The minimum values did not show a clear geographic pattern, but they never were lower in southern sites than in northern sites within the same marine basin. The general picture is that clams living in southern populations were exposed to higher temperatures during longer time periods along the year than those living in the north, as assumed in the experimental hypothesis. Average salinity values (PSU) in the three years previous to the experiment were 31.9 ± 5.4 for AN, 35.8 ± 1.7 for AS, 37.5 ± 1.9 for MN, and 37.8 ± 1.1 for MS. These values showed the expected pattern of higher salinity in the Mediterranean locations.

**Figure 2.**
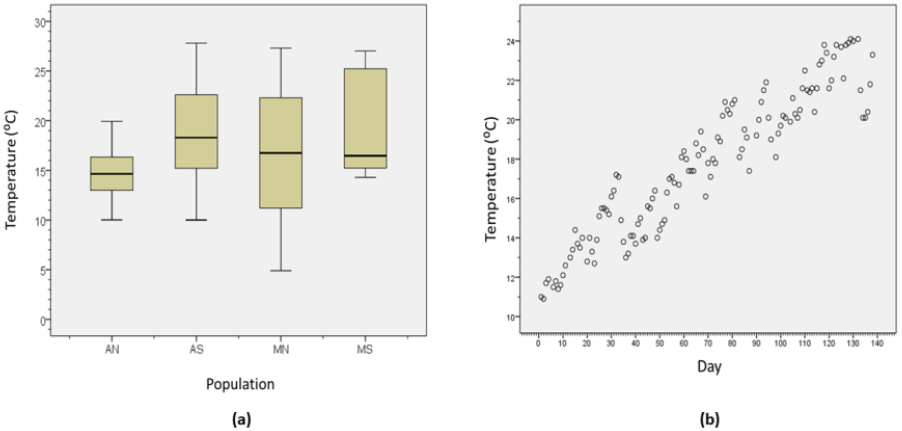
(a). Seawater temperature variation at the 4 sampled localities in the three years previous to the experiment. (b). Daily average seawater temperatures registered during the experimental period.

Temperatures of the seawater in the experimental setup increased steadily from 13.8 ⁰C to 24.1 ⁰C during the course of the experiment (Figure 2B), and fell in the range recorded in the previous 3 years in AS (Figure 2A). This indicates that the temperatures during the experiment did not depart from the usual range experienced by clams in this locality in recent times. But, on the other hand, they were higher than those experienced normally by some populations, such as AN. Salinity varied between 31.0 and 36.5 PSU with no specific temporal pattern.

### 3.2. Population genetics

Allelic frequencies for the 6 analyzed genetic markers in the four studied populations are given in Table S1 (Supplementary Information). Bayesian analysis of population structure of the 4 samples specific of this study, pooled with the 11 samples studied by [30], ascribed AN and AS samples to the Atlantic race and MN and MS to the W Mediterranean race, as expected (Figure S1, Supplementary Information). FST based on the four samples was 0.13 (P<0.001), and varied between 0.063 and 0.24 for pairwise comparisons. The neighbor joining tree based on pairwise FST separated the populations in two groups, according to their race, which reflects the higher average differentiation between the Atlantic and the Mediterranean basins, as compared to within-basin population differentiation (Figure 1, inset).

### 3.3. Gene expression variation across populations (one-way ANOVA)

Our filtering procedure rendered expression values for 11,025 probes. Results of the exploratory analysis of principal components did not reveal extraordinary departures from the general group of samples (Figure S2, Supplementary Information). Significant differences (P < 0.05) in expression level across populations were detected at 2,268 probes by the one-way ANOVA, of which 418 (3.8 %) were significant at 5% FDR (Table 1 and Supplementary Results File 1). Cluster analysis of this set of 418 probes using Euclidean genetic distances among samples showed that individuals sampled in the same locality clustered together (Figure 3), and that individuals sampled in different localities formed separate clusters. The clams from MN were the most transcriptomically divergent, and were grouped in the most basal branch of the tree. The samples from AN showed the least intrapopulation transcriptomic divergence (shortest branch lengths). Downregulated genes were more frequent than upregulated genes in all populations, except MN (Figure 4A).

**Table 1.**
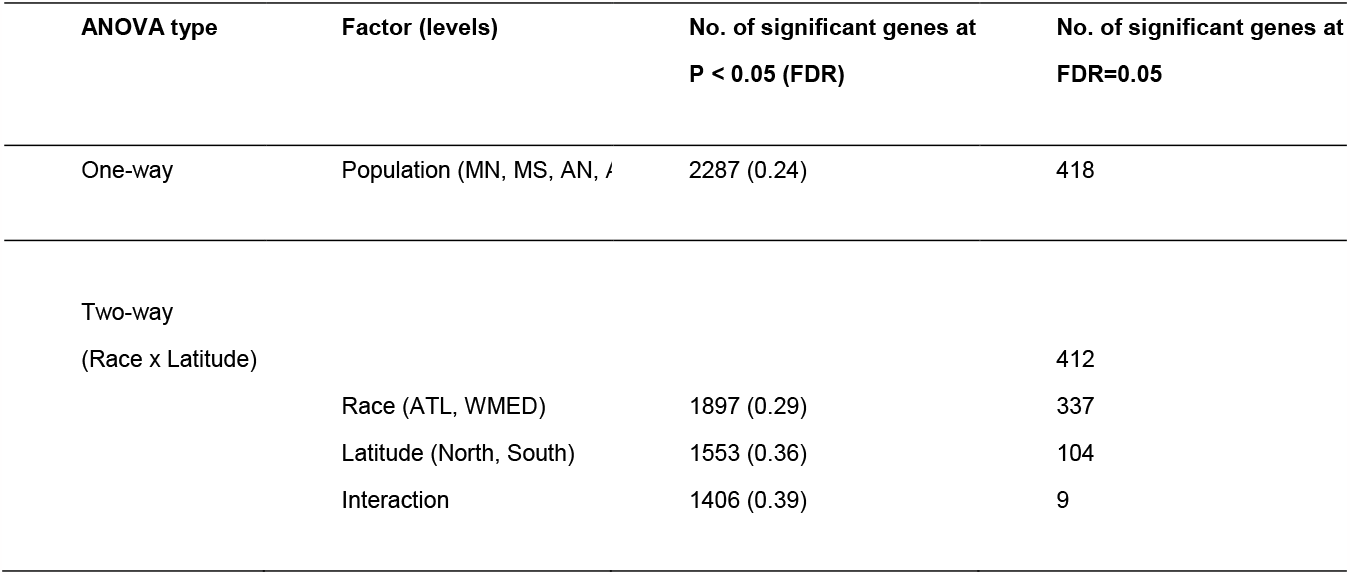
Results of ANOVA on gene expression across clam populations. FDR: false discovery rate.

| ANOVA type | Factor (levels) | No. of significant genes at<br>$P < 0.05$ (FDR) | No. of significant genes at<br>FDR=0.05 |
| --- | --- | --- | --- |
| One-way | Population (MN, MS, AN, / | 2287 (0.24) | 418 |
| Two-way<br>(Race x Latitude) |  |  | 412 |
|  | Race (ATL, WMED) | 1897 (0.29) | 337 |
|  | Latitude (North, South) | 1553 (0.36) | 104 |
|  | Interaction | 1406 (0.39) | 9 |

**Figure 3.**
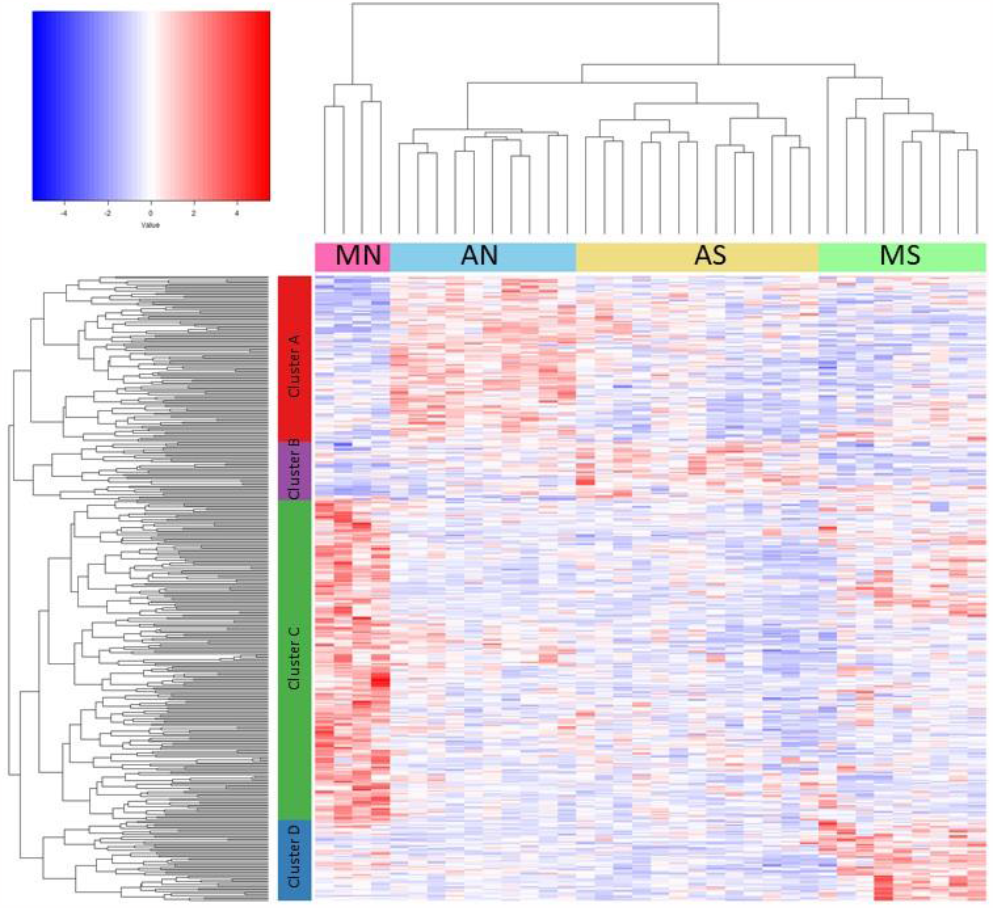
Heatmap of normalized gene expression and cluster analysis of distances between populations and genes, based on the set of genes that were significant (5% FDR) in the one-way ANOVA.

**Figure 4.**
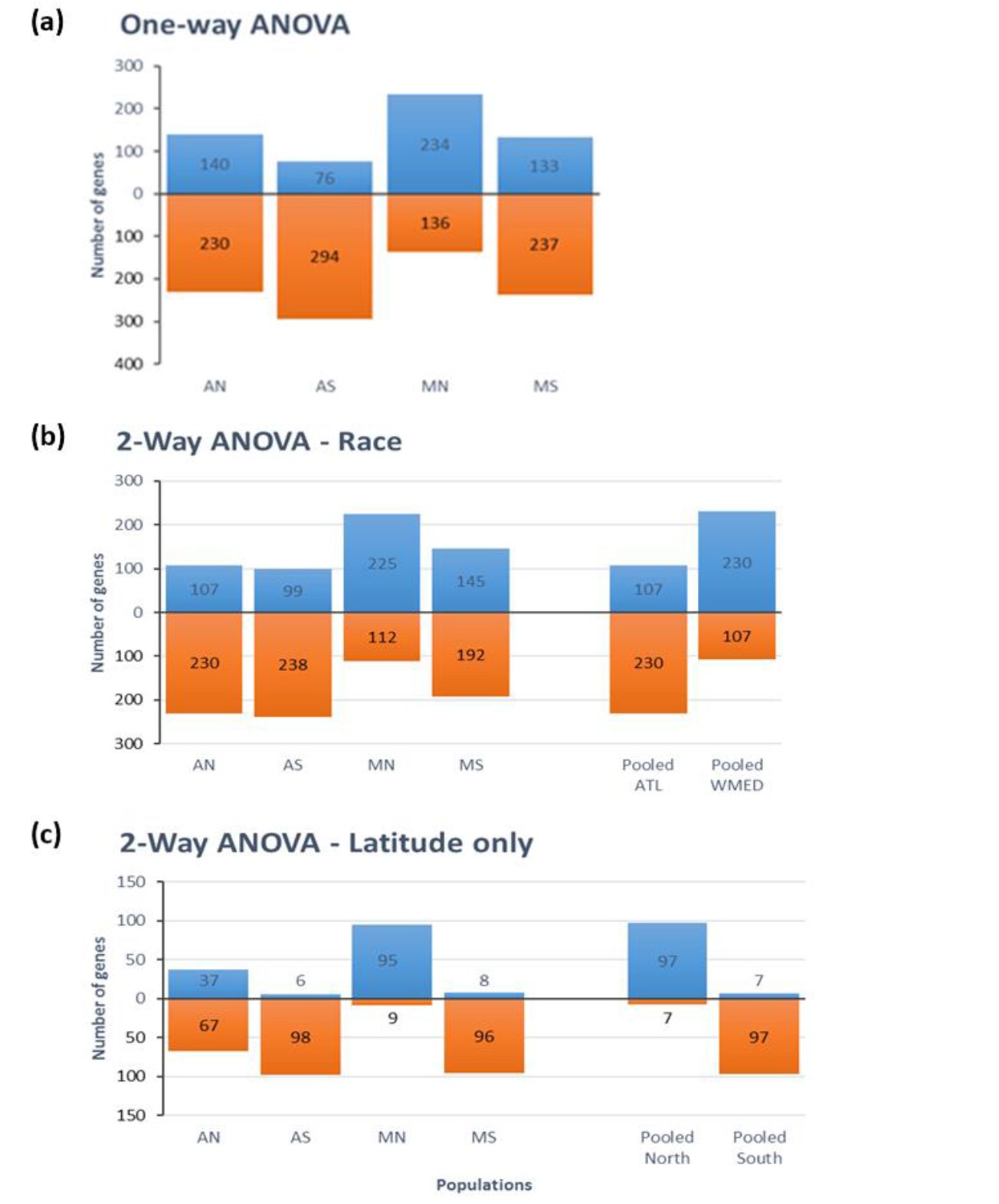
Bar charts showing the number of up-regulated (blue) and down-regulated (orange) genes according to populations and type of statistical test. Numbers for races and latitudes are also given when appropriate [(b) and (c)].

Cluster analysis of significant genes showed that populations featured characteristic expression patterns at particular groups of genes. Specifically, genes showing differential expression across populations grouped in four main gene clusters (Figure 3). Cluster A comprised 111 genes which were overexpressed mainly in AN. Enrichment tests based on their associated GO terms (Supplementary Results File 2) indicated that this cluster was enriched for genes involved in purine nucleoside metabolism and/or electron carrier activity such as cytochrome c oxidase subunit IV, methylthioadenosine phosphorylase, ATP synthase, or adenylate kinase. Cluster B comprised 54 genes which were more highly expressed in AS, and showed no significant GO term enrichment. Cluster C included 213 genes overexpressed in MN, and was enriched for genes related to ribosomal activity, peptide biosynthesis and glutathione metabolism. Finally, cluster D comprised 39 genes which were overexpressed in MS, and was enriched for genes related to cellular respiration such as cytochrome c oxidases and NADH dehydrogenases.

### 3.4. Gene expression differences associated to race and latitude (2-way ANOVA)

Results of the 2-way ANOVA (Race x Latitude) are shown in Table 1. This analysis detected significant differences (5% FDR) at 413 probes, of which 278 probes (50%) had been significant in the 1-way ANOVA. The genes significant in the 2-way ANOVA included 337 probes significant for Race (87%), 104 for Latitude (25%) and 9 for their interaction (2%). Some probes were significant for the two factors, or for a combination of factors and interaction (Figure 5A). Specifically, twenty-nine probes which showed significant signal differences between races also showed significant differences for latitude. Similarly, 6 probes that exhibited significant Race x Latitude interaction also showed significant effects for one of the two factors (3 probes), or both (3 probes). In total, 35 out the 413 significant probes were significant for some combination of race, latitude and interaction (Figure 5A). The 337 probes which were significant for the Race factor in the 2-way ANOVA (Figure 5A) are candidates for expression differentiation caused by the genetic divergence that occurred during one or more episodes of Atlantic-Mediterranean population subdivision in the Pleistocene. In this group of genes, average signal intensity was higher in the Atlantic than in the Mediterranean populations for 107 probes (32%), and higher in the Mediterranean populations for 230 probes (68%) (Figure 4B).

**Figure 5.**
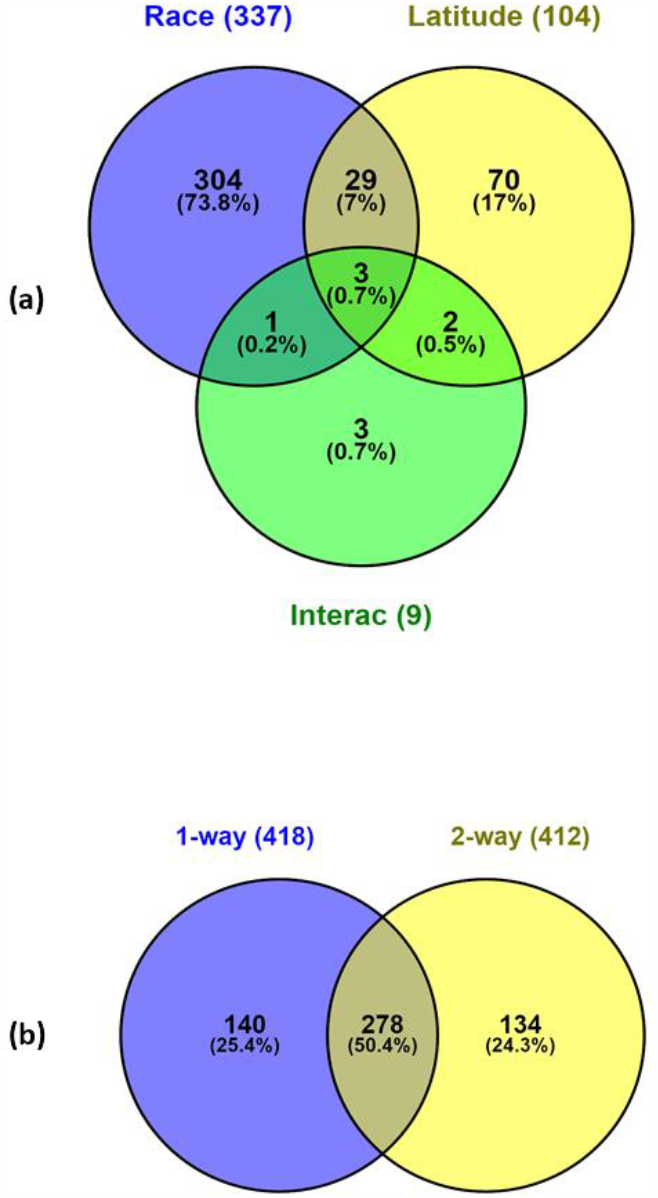
Venn diagrams showing the number of DEGS (FDR = 0.05) of gene expression data. **(a)** 2-way ANOVA (Race x Latitude). **(b)** Comparison of the DEGs between 1-way and 2-way ANOVA.

GO-terms could be obtained for 173 race-significant genes by BLAST against the Swissprot database and for 200 through BLAST against the *Danio rerio* genome (Supplementary Results File 1). The list of GO-terms in the Cell Component category indicate that DEGs code for proteins located in the cytoplasm, the nucleus, the membranes and the extracellular space. Within the cytoplasm, DEGs coded mainly for proteins of the cytosol, the cytoskeleton, the mitochondria, the Golgi system, with other parts in lesser numbers. This diversity of cell locations runs parallel to a large number of GO terms (169) associated to a very diverse set of functions in the Biological Process category, the most frequent being Translation and other functionally related terms, proteolysis, oxidation-reduction, regulation of transcription, transport and metabolic processes. Fisher exact tests indicated overrepresentation of 44 GO terms at P<0.05 in this set of genes, although this term collection is highly redundant (Figure 6). The term that grouped the highest number of probes was Protein metabolic process (46 probes). Nineteen additional overrepresented terms were related to protein synthesis and ribosomal function, comprising 22-34 probes. The remaining overrepresented terms were related to cell homeostasis, development, differentiation, carbohydrate metabolism, extracellular part, peptidase activity and immune system (in decreasing number of probes). The KEGG pathway dre03010:Ribosome was also enriched. The heatmap in Figure S3 (Supplementary Information) shows the between-race variation for the subset of 105 genes associated to the significantly enriched GO terms. These genes include a large number of ribosomal proteins and other proteins related to protein biosynthesis, of which 14 were more highly expressed in the Mediterranean populations and 7 in the Atlantic. Five protease DEGs (cathepsin and other proteases) were more highly expressed in the Mediterranean and only 1 in the Atlantic. The five DEG coding for proteins of the mitochondrial electron transport chain (cytochrome oxidases, NAD dehydrogenases) and the functionally related ATP-synthase F1 subunit were more highly expressed in the Atlantic.

**Figure 6.**
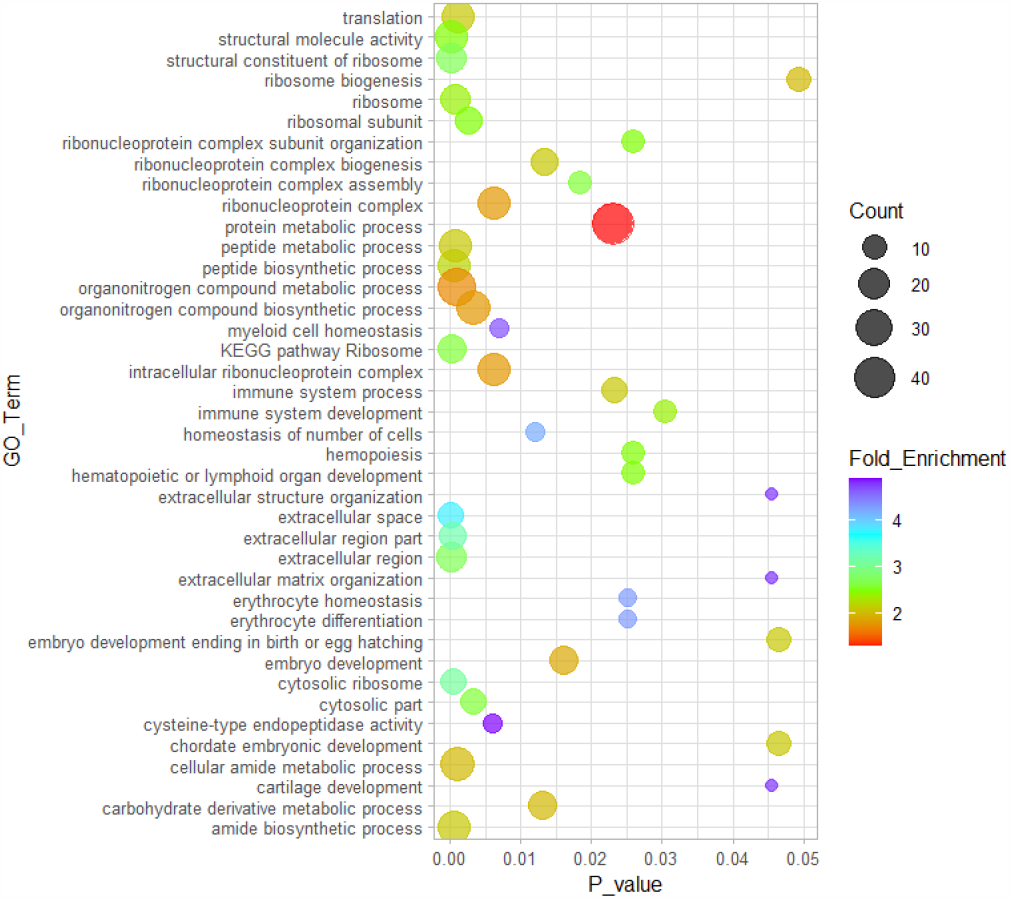
Overrepresented Biological Process GO terms (P <0.05) in the set of genes which were significant for the Race factor in the 2-way ANOVA.

Out of the 104 genes which were significant for latitude in the 2-way ANOVA, 32 were also significant for Race and two were significant for the Race x Latitude interaction (Figure 5A). The remaining 70 genes, which showed significant transcriptomic differentiation for latitude but not for race, are candidates for adaptive gene expression variation related to temperature differences between northern and southern populations. We will refer to them as the “latitude-only” gene set. The average signal was higher in the northern populations for 63 probes (88 %) in this gene set, and higher in the southern samples for only 7 probes (Figure 4C). When the latitudinal expression change was considered separately within each race (i.e.: AN compared with AS, and MN with MS), this pattern remained the same, with only one exception (gene #4479, with no BLAST hit; File S1 in Supplementary Results File 1). These results indicate that after transplantation to the warm waters of southern Portugal, the clams which were native to the northern, cool environments had higher transcription rate than the clams native to the southern environments for most of those genes. We found 13 GO terms overrepresented in this group of genes at P<0.10 (Figure 7). The term Cellular processes involved the largest number of genes (24), with an average fold enrichment of 1.19X. The highest enrichments were found for the terms transferase activity and glutathione transferase activity (23X and 27X, respectively). Three other enriched terms (GO:1901564, GO:0006790, GO:0043603) included genes coding for enzymes of glutathione metabolism. The three KEGG pathways which were significantly enriched in the “latitude only” gene set were drug metabolism – cytochrome P4502, metabolism of xenobiotics by cytochrome P450, and glutathione metabolism. The genes responsible for the significance of these three pathways gave BLAST hits to glutathione S-transferase (GST) omega (two cases) and GST-rho of zebrafish (genes #3702, #12815, and #8274 in Supplementary Results File 1). The three genes were upregulated in the northern populations exposed to southern temperatures (Figure S4, Supplementary Information). The results of BLASTX against data bases other than Ensemble zebrafish revealed another two genes in the latitude-only gene set (genes #8304 and #4170, Supplementary Results File 1) with hits on GSTs of other species. Another gene present in this gene set codes for glutathione synthetase, an essential protein for the provisioning of reduced glutathione in the cell (gene #4887, Supplementary Results File S1). All these results underscore the importance of upregulation of glutathione-related genes in northern populations.

**Figure 7.**
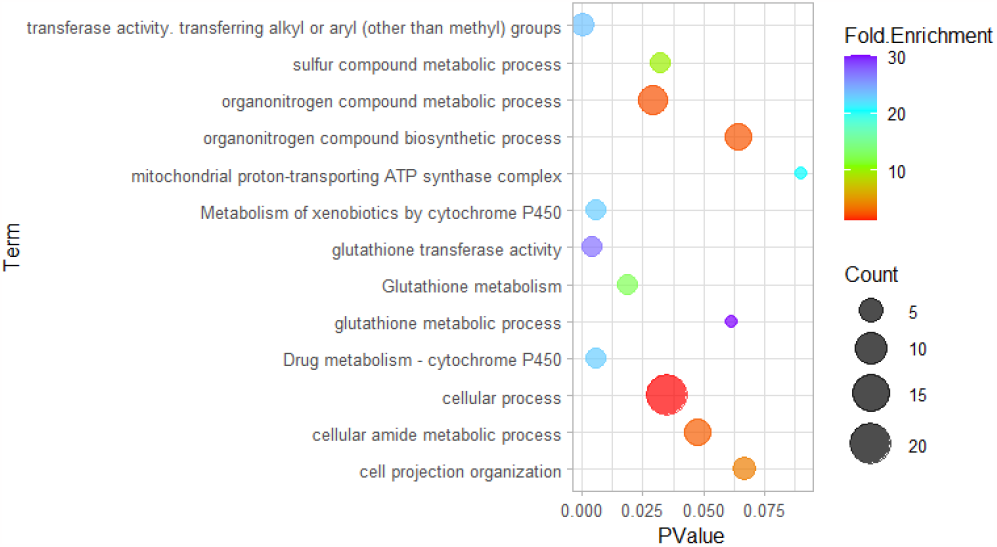
Overrepresented Biological Process GO terms (P <0.05) in the set of genes which were significant only for Latitude in the 2-way ANOVA.

The GO term mitochondrial proton-transporting ATP synthase complex showed also P-value <0.10 and a very high fold enrichment (20X) in the latitude-only gene set (Figure 7). This term was associated with two genes coding for subunits of the FoF1-ATP-synthase enzyme (ATPK) (genes #5665 and #7588 in Supplementary Results File 1, and Figure S4). Two other genes in the “latitude only” gene set, but showing no high-confidence BLAST against any zebrafish gene, and therefore missed in the GO term overrepresentation tests, seemed to correspond to ATPK as well, as indicated by their similarity to the Drosophila melanogaster ATPK in the SwissProt database (genes #1210 and #10018 in Supplementary Results File 1). Their presence in the latitude only gene set gives additional support to the overrepresentation of ATP synthase complex in this group of genes.

Nine probes gave significant results for the Race x Latitude interaction (Table 1). The patterns of variation at these probes showed a clear distinctiveness of the MN population. A set of seven probes showed higher average signal in MN (Figure S5, Supplementary Information), of which five showed no BLASTX match, and three corresponded to genes coding for spermatogenesis-associated protein 1 (SPAT-1), stress response protein nhaX and protein-L-isoaspartate(D-aspartate) O-methyltransferase (PCMT-1). Another two genes showed the lowest expression value at MN, and corresponded to T-complex protein 1 subunit alpha, Nucleolar complex protein 3 homolog, prohibitin-2 (2 probes) and Eukaryotic translation initiation factor 2-alpha kinase 4.

### 3.5. Test of selection on gene expression variation

The results of EVE selection test are reported for each gene in the Supplementary Results File 1. For the null hypothesis of higher variance among populations than predicted by the phylogeny, tests were significant at P< 0.05 for 235 genes. However, none were significant after multiple test correction by the BH procedure (P > 0.10 in all cases).

## 4. Discussion

Taking together the results of the one-way and two-way ANOVAs, 553 genes showed differences in gene expression among localities after transplantation of clams to the experimental facility and gradual acclimation to a common environment for a period of ca. 3 months. This is 5% of the genes included in the microarray. The number of DEGs has been estimated by applying a significance threshold of 5% FDR to account for multiple testing, which is quite a stringent threshold. Gene expression differences probably occur in a higher number of genes. The number of differentially expressed genes in our study at a single-test significance threshold of P < 0.05 was 2287 (20%), and this can be considered well above the real number, since multiple testing would result in a large fraction of false positives. Therefore, we can conclude that an important number of genes (at least 5%, but less than 20%) showed differences in expression across populations of R. decussatus clams. Previous studies in animals have reported highly variable fractions of genes showing differences in gene expression among populations: 0.1 % in a marine snail [46], 7.8% in humans [47], or 12% in the killifish [48], to cite just a few diverse examples. These studies are only loosely comparable, since they differ in critical aspects such as the technique used (microarray or RNA-seq), the number of replicates, the sample sizes, the statistical significance threshold selected, and on whether the organisms had been acclimated in the laboratory before performing the study or they were analyzed immediately after they were sampled in the field. Thus, the number of DEGs found among populations of GCS clam is within the range of values observed in other animal species.

The 1-way and the 2-way ANOVA rendered similar numbers of DEGs among populations (418 and 412 genes, respectively). The similarity of these numbers hides the fact that only 278 probes (50%) were significant in both analyses (Figure 5B). 140 genes (24%) that were significant in the 1-way ANOVA were not significant in the 2-way ANOVA. This suggests that, although the factors race and latitude were capturing a substantial amount of the interindividual variation in gene expression, the variation at one quarter of the DEGs could be due to other factors. Conversely, 134 probes which were significant in the 2-way ANOVA were not in the 1-way ANOVA, suggesting that the partition of the variance performed by the 2-way ANOVA resulted in a statistical power increase that led to detecting many new DEGs.

Our study was based on a “common garden” design, addressed to detect genes whose expression at the population level were potentially affected by genetic differences between populations. Under the logic of this design, the observed gene expression differences between clams sampled in different localities, but kept under a common environment for a long time, are interpreted as innate differences in gene expression. There are two potential sources of innate differences. One is the existence of differences in the allelic composition of the populations at the genomic regions responsible for transcription regulation. These genetic differences among populations would have been produced either by the classic neutral population genetic mechanisms of mutation, genetic drift and gene flow limitation, or by natural selection. The second potential source of innate differences is the action of epigenetic memory mechanisms elicited by the local native environments, which could be still acting after the acclimation to the experimental setting [49]. For example, differences among populations in DNA methylation patterns potentially involved in epigenetic regulation have been shown in in another bivalve, the oyster *Crassostrea gigas* [50,51]. Specific studies will be necessary to determine the existence and extent of epigenetic differences influencing the transcription patterns associated to the response to seawater warming and those observed between the two clam races.

Irrespective of whether epigenetic factors could be playing a role, the most obvious source of innate transcriptional differences in the studied clams is genetic drift resulting in genetic differentiation of transcriptional regulatory genomic regions. Genetic drift should have had a principal role, as the two races involved in this study appeared as a result of the species subdivision in the Pleistocene [30]. Moreover, present day limitation to gene flow at the Almeria-Oran front probably contribute to maintaining the genetic differences between the two races as well, as it seems to happen in other species (e.g.: [30,52]. An additional source of transcriptional differentiation due to genetics is isolation by distance between populations within the Atlantic and the Mediterranean marine basins. Previous studies with microsatellites showed that the amount of genetic differentiation (*F*_*ST*_) among pairs of Atlantic populations reached 0.08, and values for two West Mediterranean populations on the Spanish coast reached 0.04 [31]. These values indicate that gene flow within each basin is relatively limited, which allows for an important effect of genetic drift at intermediate to high geographic distances in the long term.

If the expression differences among populations were the result of genetic drift alone, the pattern of overall gene expression differences among populations should match the pattern of genetic relationships derived from the population genetic data, as they appear in the phylogenetic tree shown in Figure 1. In other words, the two Atlantic and the two Mediterranean populations should be transcriptionally more similar to each other than to any population of the other race. The cluster analysis of the DEGs detected by the 1-way ANOVA departs slightly from this expectation. Although the samples from the two Atlantic populations clustered together, the MN population appeared basal to the other 3, suggesting that this population was transcriptomically more differentiated (Figure 3). Results of other analyses also indicate some differences at MN. For example, contrary to the other 3 populations, the MN population showed a higher proportion of up-regulated than down-regulated genes (Figure 4). Moreover, genes showing significant interactions in the 2-way ANOVA showed a clearly distinct level of expression in the MN population (Figure S5, Supplementary Information). These observations suggest that factors other than the historical population split and isolation by distance have influenced the pattern of gene expression differences across clam populations observed in this study, and that those factors are, at least in part, specific of MN. The clams collected in that population experienced a high mortality since the very moment of its capture in the lagoon of Thau, while mortality in the clams collected in the other localities was very low. This means that the health status of MN clams was compromised already at the moment of sampling. We can only speculate about the causes of such health impairment. Harsh winter conditions in the lagoon could have had a role, as the temperatures recorded in the lagoon in winter were much lower than in any other population (Figure 2), and we have observed winter mortality of clams maintained in captivity at the IATS-CSIC facilities due to similarly low seawater temperatures while performing other studies (unpublished results). The effect of a disease seems less probable, as there were not significantly enriched GO terms related to disease response in the MN-specific group of DEG. Alternatively, the pattern of transcriptomic differentiation of MN population could be due to a different genetic makeup of the population. Some support for this explanation comes from the population genetic data. Allele C at locus TBP has been recorded previously in clam populations from the Adriatic and Aegean seas only [30], but in the present study it appeared in MN at a very low frequency (Table S1, Supplementary Table and Figures). This points to the presence of some individuals of Eastern Mediterranean ancestry among the MN clams, which could be native or introduced. More detailed genetic studies of the clam populations in south-east France are necessary to clarify this point.

In spite of the particularities detected in the transcription profiles at the MN population, the results point to a large fraction of DEGs not being affected by these local features. Actually, the results showed that the observed patterns of transcription variation among clams at three quarters (413) of the 552 detected DEGs can be explained by two single factors: the race the clams belong to and the latitude of the locality where they were sampled. Ninety per cent (373 out of 413) of the significant genes in the two-way ANOVA gave significant results for only one of the two factors, and only 38 genes showed significant variation for both factors and/or their interactions. Overall, the race factor seems to have had a stronger effect on transcriptomic differentiation among populations, as it explained the variation observed at 337 genes (62 % of the 552 detected DEGs), while the sampling latitude explained only one third of that (104 genes, or 19 %).

### 4.1. Transcriptional differences between races

Previous work using intron polymorphisms and microsatellites showed that populations of GCS clam from the Atlantic and the western Mediterranean were differentiated in allele frequencies at a large fraction of the scored genes, and therefore can be considered different races, which were called ATL and WMED, respectively [30,31]. A third race comprising populations from the Aegean and Adriatic seas (AEGAD) was also detected [30], but populations of this race have not been included in this study. Our initial hypothesis was that population subdivision should have resulted in the accumulation of nucleotide differences along the whole genome, including regulatory regions of coding genes, and therefore should result in differences across populations in the transcription levels of many genes The results of the 2-way ANOVA showed statistically significant expression differences between Atlantic and West Mediterranean populations for 337 genes (3%), confirming the previous hypothesis. This result constitutes the first transcriptomic characterization of the two clam races, and also (to our knowledge) of any marine species exhibiting genetic or phylogeographic differentiation across the Mediterranean Sea and the nearby Atlantic Ocean. Although other studies of population transcriptomics have included samples from regions separated by phylogenetic breaks, specific characterization across the breaks has not been properly addressed in those studies. Increased transcriptomic differentiation between Atlantic and one single Mediterranean population separated by a phylogeographic barrier was observed in the seagrass *Zostera marina* [24]. However, to our knowledge no other transcriptomic screening addressed specifically to a phylogeographic system has taken place so far.

The results cast several questions regarding the clam races. One of the most interesting questions is: to what extent do the observed differences in transcription result in differences in the physiology? We have approached this question by performing the functional characterization of the differentially expressed transcripts with the Gene Ontology (GO) system, and by enrichment tests of GO categories. This approach has several limitations, the most important being that GO annotation in bivalve mollusks (as well as non-model invertebrates in general) is quite underdeveloped. Only 40% of the DNA sequences used to design the microarray probes could be associated to a known coding sequence in other organisms by BLAST family tools, and only a fraction of them had associated GO terms [38]. It is also well known that the transcript levels are only loosely related to the amount of protein in the cells [53], so transcription differences between individuals or groups are only indicative of potential physiological differences. For all these reasons, the conclusions based on transcription differences and GO analyses should be taken with great care and will require independent testing in other clam samples by additional studies [54].

If transcriptional differences would have been generated by random genome-wide variation resulting from mutation and genetic drift after the Pleistocene split of the species in Atlantic and Mediterranean subpopulations, it would be expected that many genes scattered over the whole genome and with different functions would show expression differences. Our results fit this expectation. GO-terms associated to the race-related set of DEGs show that this gene set was functionally very diverse, with proteins pertaining to nuclear, cytoplasmic, membrane, cytosolic and extracellular parts, and showing a large variety of molecular functions.

Similarly, random differentiation would result in ca. 50% upregulated genes in each marine basin. However, 68% upregulated genes were observed in the Mediterranean, which departs substantially (although not dramatically) from this expectation. GO-term enrichment tests indicate that some GO categories were significantly overrepresented in the set of race–associated DEGs. The most outstanding were a group of GO terms associated to a total of 45 genes coding for proteins involved in the translation machinery (Supplementary Results File 1). Nineteen of these genes code for ribosomal proteins, and most of them (14) were upregulated in the Mediterranean race. Other genes involved in translation were eukaryotic translation factor 3 and NSA2 ribosome biogenesis homolog. These results suggest that protein synthesis routes could show important differences in the two races, although it should be considered that the regulation of expression of ribosomal proteins is only shallowly understood in metazoans. Moreover, these proteins may have other functions aside from their role in the ribosome assembly [55].

In addition to GO terms related to protein biosynthesis another 18 terms were significantly enriched in the race-associated DEGs, which can be summarized in three categories: development, cell differentiation and extracellular space. These terms are associated to some of the ribosomal proteins and translation related proteins, clearly showing the large influence that the differences in the transcription levels of ribosomal and protein biosynthesis-related proteins between the clam races might have on other aspects of their biology aside of protein synthesis. However, these terms include many other proteins as well. Especially interesting from the point of view of physiology is the significant enrichment for carbohydrate derivative metabolic process. This term involves 15 DEGs, some of which code for proteins of the mitochondrial energy production processes, such as adenylate kinases, cytochrome c oxidase subunit IV, ATP synthase subunits, or NADH dehydrogenase. Other DEGs code for enzymes involved in the post-translational modification of proteins, such as glycosilations and aminations (glutamine-fructose-6-phosphate transaminase 1), and others play a role in the biosynthesis and transport of glycosylated lipids (phosphatidylinositol glycan anchor biosynthesis, class X, and GM2 ganglioside activator). GO-term enrichment tests also show enrichment for terms related to proteins of the extracellular spaces. Six DEGs induced significance of the cysteine-type endopeptidase activity GO term. Cysteine type endopeptidases have a role in catabolism of proteins in the lysosomes. These proteins have important roles in many aspects of the cell biology such as tissue growth, cell differentiation, signaling, or immunity. The GO terms and genes summarized above are just a fraction of the whole set of GO terms associated to genes that are significantly differentiated in expression between the two races. The large number of enriched GO terms suggest that many cell and physiological functions can be affected by differential transcription levels between the two clam races and, therefore, might result in physiological differences between them. Physiological differences between clam races is a topic that should be examined in detail in future studies.

The great majority of observed expression differences between races should be the result of the genomic differences accumulated during the separation of the Atlantic and Mediterranean lineages during the Pleistocene due to genetic drift. However, we cannot exclude that environmental changes that took place during the glacial periods had influenced the transcription rates of clams in their Atlantic and Mediterranean refuges. The fact that a fraction of the genes that have been found significant for the Race factor in the 2-way ANOVA were also significant for the latitude factor suggest that they may be also responding to the temperature differences between northern and southern localities in present times, but this response could have been modelled by differences in temperature between the two glacial refuges (see next section).

### 4.2. Transcriptional differences associated to latitude

Northern clams showed increased transcription of 104 probes, as compared with southern clams, in the 2-way ANOVA. As many as 34 genes were also significant for race, but 70 were significant for latitude only, and they are candidates for transcriptional response to temperature change. GO term enrichment tests showed clear overrepresentation of two types of enzymes: glutathione transferases (GST) and ATP synthase subunits (ATPK). GST is a family of enzymes which is well known for their participation in glutathione-based xenobiotic detoxification phase II mechanisms [56–58]. They also neutralize secondary metabolites produced during oxidative stress [57]. Temperature increase is a source of oxidative stress and, accordingly, several studies have seen GST expression related to increased environmental temperature in the gills of marine bivalves [59,60]. Some authors have indicated that temperatures of 25ºC or higher are stressful for the GCS clam [61]. This temperature was approached, but never reached in the present study. Lack of differential stress experienced by northern and southern clams is also suggested by the absence of heat shock proteins in the group of DEGs. Therefore, it is probable that northern clams have simply higher transcriptional levels, either constitutively or as a plastic response to temperatures that are less frequent in their normal habitat. In addition to the enrichment of GST, other data support the involvement of glutathione-based xenobiotic detoxification phase II mechanisms. Specifically, two other probes with BLAST hits to enzymes related to glutathione metabolism were significant for the latitude factor: glutathione peroxidase (gene #1975) and glutathione synthase (gene #4887)(Supplementary Results File 1).

ATP-synthase (ATPK) is a multimeric enzyme with the essential role of coupling the electron transport chain with the synthesis of ATP in the cellular respiration process inside the mitochondria [62]. Due to its position in the proton transfer enzyme network, it also has a role in protecting towards oxidative stress. Several studies have shown a relationship between ATPK expression and environmental temperature in aquatic organisms, including bivalves. Increased expression of ATP synthase has been found in the muscle of cold-acclimated carps (*Cyprinus carpio*) [63,64]. Li et al. [65] reported increased expression of ATPK at both high and low stressful temperatures in the razor clam *Sinonovacula constricta*. In the mussel *Geukensia demissa*, ATPK increased expression in autumn, in parallel to a decrease in water temperature [66]. Wijers et al. [67] found increased amounts of ATPK in human skeletal muscle cells taken from biopsies of human males exposed to cold temperature during three days. All these studies suggest that expression of ATPK is frequently upregulated in response to a decrease of environmental temperature. These studies support the view of the observed higher expression of ATPK in northern clams is a result of constitutive expression associated to their cool native environment.

GST and ATPK represent only a few of the 107 genes that have shown differential expression between northern and southern clams. While only enriched terms related to oxidative stress response and ATPK were found in the “latitude-only” set of genes, it is worth noting the presence in it of several genes coding for proteins of the mitochondrial electron transport, such as NADH dehydrogenases or cytochrome b, which interact intimately with ATPK, and were upregulated in the northern populations. All together, these observations strongly suggest that upregulation of some electron transport chain proteins is constitutive in the GCS clams living in cool environments. This conclusion is line with studies of Antarctic fishes, in which upregulation of oxidative chain genes has been described and explained as an adaptation to the cold environment [68].

A fraction of the genes which have been found significant for the Latitude factor in the 2-way ANOVA were also significant for the Race factor, indicating expression differences between races. It is possible that these genes are responding to the temperature differences between northern and southern localities in present times, and this response could have been modelled in the past by differences in temperature between the two glacial refuges. This possibility makes sense in the light of the different opportunity of the clam populations to move latitudinally during glaciations. Movement of Mediterranean populations from north to south in response to the advance of the cold wave southwards would have been restricted by the presence of the northern African coasts. However, the populations in the Atlantic could have moved more to the South, along the west coast of Africa, staying in their optimal, warmer environment. These differences could have resulted in natural selection operating at the Mediterranean population to cope with the cooler environment, favoring the action of natural selection to modulate transcriptional levels at genes related to thermal response. These differences could have remained until present times giving rise to our observations.

### 4.3. Potential adaptive role of the observed differential expression

A recurrent topic of research regarding transcriptomic variation among individuals and populations is whether the differential expression is caused by the accumulation of neutral polymorphisms in the genome, or they are due to adaptive changes in the DNA driven by natural selection. Several ways to perform tests for adaptive transcription differences have been proposed, but all seem to have pitfalls [69]. Previous studies of gene expression differences among conspecific populations have used comparisons between DNA and quantitative genetic variation with F_ST_/Q_ST_ ratios [70], phylogenetic contrasts [48] and, more recently, partition of variance [45]. Variance contrast tests were developed to deal with study designs like the one used in this work, based in population sampling. In this study, we used this approach using the EVE (Expression Variance and Evolution) software [45]. We found no clear evidence for selection, because even though more than 200 genes were significant for the test, correcting for multiple testing resulted in no gene showing significant deviation from the neutral expectation. This result should be taken with great care for several reasons. Firstly, the test is based on the partition of phenotypic variance according to the phylogeny assuming a complete split of taxa, [45], and the effect of gene flow among taxa is not considered. Moreover, even if the tests were robust with regard to deviations from the model assumptions, the power of these tests is probably low in this particular case because of the small number of populations scored. We know of only one application of EVE to intraspecific populations, in which significant results were obtained [71]. That study differed from ours in three ways: slightly larger number of populations, use of RNA-seq instead of a microarray, and the fact that the samples compared were not kept in a common environment before the expression study, and therefore were influenced by environmental differences among the sampled habitats which probably resulted in an increased gene expression variation due to plasticity, which could confound with the genetically-based variation.

In the absence of powerful neutrality tests, correlations with environmental parameters provide an alternative way to look for candidates to adaptive evolution. Our factorial sampling scheme (race x latitude) allowed to separate the effects of genetic differentiation between races from the effects of the latitudinal thermal gradients operating in the study area. The set of 70 genes that showed latitude related differential expression, but not expression differences between races, constitute a set of candidate genes for response to thermal variation. As these latitude-associated differences have been observed after a long period of acclimation to a common environment, the conclusion is that they are caused by genetic differences in the expression of the genes, which have been driven by adaptation to the thermal characteristics (and possibly other factors associated to temperature) of the northern habitats where the clams were sampled.

An alternative explanation for the observed latitude-related differences in gene expression is that they are due to neutral genetic differences between the north and south populations in each basin. Molecular dating suggests that the differentiation between the Atlantic and West Mediterranean races was due to historical separation in the Pleistocene, not earlier than 400.000 years before present (YBP) and as far as 2 million YBP [30]. Therefore, the clam populations of the two basins have been evolving independently during a long period of time in the past, resulting in highly differentiated races. Moreover, present-day gene flow is probably limited between them due to the Almeria-Oran Oceanografic Front (AOF) barrier. As a consequence of this independent evolution in the two basins, it would be expected that, just by chance, many genes that show gene expression differences between north and south populations in one basin, would show differences in the opposite direction in the other basin. However, the observation of only 6 out of 70 genes showing contrasting patterns of expression between north and south in the two basins argues against this possibility, and supports an adaptive role of the observed latitude-related expression variation in the two parallel thermal gradients that occur at both sides of the Iberian Peninsula.

## 5. Conclusions

We have shown that populations of *R. decussatus* that constitute the two races separated by the Atlantic-West Med phylogeographic break, and spanning parallel latitudinal gradients in temperature, show clear differences in gene expression at a remarkable portion of their genes after a relatively long period of acclimation to a warm, Atlantic environment. A large fraction (3/4) of these differences were explained by simple, direct effects of two factors: the race the clams belonged to and the latitude where they were sampled. The differentiation patterns were different for each of these causal factors. In the case of racial expression differences, most genes showed higher expression in the Mediterranean, but a significant fraction (40%) showed the opposite pattern. This observation is concordant with an explanation based on random differentiation of populations due to genetic drift during historical isolation caused by the Pleistocene sea level changes, although additional implication of natural selection cannot be discarded. An important fraction of the DEGs between races are related to the protein biosynthesis machinery and ribosomal function, which suggest that the two races could show important differences in their functional aspects regarding adaptation to global warming and aquaculture. Additional research will be required to find out if the observed differences between races are a response of the Mediterranean race to the transplantation to Atlantic waters, or were already present in their native habitats. In the case of the genes significant for latitude, the observation of higher expression at almost all significant genes in clams coming from the northern localities, which usually experience lower temperatures than those in the south, independently of their Atlantic o Mediterranean genetic background, suggest that we have recorded a constitutive expression response to the N-S temperature gradient that may have a genetic basis. The analysis of GO term enrichment in the set of genes whose expression was correlated with latitude uncovered that glutathione metabolism and energy production at the mitochondria are key metabolic routes that should be explored in the quest to understand the adaptation of grooved carpet-shell clams, and possibly bivalves in general, to ocean warming.

## Supporting information

Supplementary Results

## Supplementary Materials

The following two files contain supporting information:

1. Supplementary Tables and Figures: Table S1 and Figures S1 –S5 (PDF format)
2. Supplementary Results File (Excel, csv format)

## Author Contributions

Conceptualization, C.S.; methodology, C.S., M.M, R.L., D.C. and L.B; software, C.S., M.M. and D.C.; validation, C.S., M.M., D.C. and L.B.; formal analysis, C.S., M.M. and D.C.; investigation, C.S., M.M. and D.C.; resources, C.S., M.M., R.L., L.C., T.P. and L.B.; data curation, C.S., M.M. and D.C.; writing—original draft preparation, C.S.; writing—review and editing, C.S, M.M., R.L., D.C., T.P. and L.B.; visualization, C.S. and D.C.; supervision, C.S. and L.B.; project administration, C.S., L.C. and L.B..; funding acquisition, C.S., L.C., T.P. and L.B. All authors have read and agreed to the published version of the manuscript.

## Acknowledgments

This work was funded mainly by grant AGL2010-16743 from the Spanish Ministry of Science and Innovation to C.S, and by funds provided by the University of Padova. Parts of the research were carried out during a stay of D.C. and C.S. at the Centro de Ciencias Marinhas do Algarve in Faro, and a stay of C.S. at the University of Padua, which were financed by the UE funded ASSEMBLE program and a BEST11 travel grant from the Generalitat Valenciana to CS, respectively. We would like to thank the following people for their help in clam collection: François Bonhomme (CNRS and University of Montpellier), Liliana Solís (Confraría de Pescadores San Bartolomé de Noia), Benjamín García (IMIDA), Michel Poitou (Station Marine de Sète-OREME), and Nicolas Bierne (CNRS and University of Montpellier). We thank Serena Ferraresso and Mariana Pauletto for their help with microarray hybridizations, and Valerie Derolez (IFREMER) and Nicolas Bierne for providing environmental data from the Thau lagoon.

## References

1. Franks, S.J.; Hoffmann, A.A. Genetics of Climate Change Adaptation. Annu. Rev. Genet. 2012, 46, 185–208, doi:10.1146/annurev-genet-110711-155511.

2. Froehlich, H.E.; Gentry, R.R.; Halpern, B.S. Global Change in Marine Aquaculture Production Potential under Climate Change. Nat. Ecol. Evol. 2018, 2, 1745–1750, doi:10.1038/s41559-018-0669-1.

3. Frölicher, T.L.; Fischer, E.M.; Gruber, N. Marine Heatwaves under Global Warming. Nature 2018, 560, 360–364, doi:10.1038/s41586-018-0383-9.

4. Somero, G.N. The Physiology of Global Change: Linking Patterns to Mechanisms. Annu. Rev. Mar. Sci. Vol 4 2012, 4, 39–61, doi:DOI 10.1146/annurev-marine-120710-100935.

5. Deliberto, A.N.; Drown, M.K.; Ehrlich, M.A.; Oleksiak, M.F.; Crawford, D.L. Feeling the Heat : Variation in Thermal Sensitivity within and among Populations. J. Exp. Biol. 2022, jeb.244831, doi:https://doi.org/10.1242/jeb.244831.

6. Batta-Lona, P.G.; Maas, A.E.; O’Neill, R.J.; Wiebe, P.H.; Bucklin, A. Transcriptomic Profiles of Spring and Summer Populations of the Southern Ocean Salp, Salpa thompsoni, in the Western Antarctic Peninsula Region. Polar Biol. 2017, 40, 1261–1276, doi:10.1007/s00300-016-2051-6.

7. Whitehead, A.; Crawford, D.L.; Whitehead, A.; Crawford, D.L. Neutral and Adaptive Variation in Gene Expression. 2007, 103, doi:10.1073/pnas.0507648103.

8. Coscia, I.; Vogiatzi, E.; Kotoulas, G.; Tsigenopoulos, C.; Mariani, S. Exploring Neutral and Adaptive Processes in Expanding Populations of Gilthead Sea Bream, Sparus aurata L., in the North-East Atlantic. Heredity (Edinb). 2011, 108, 537–546, doi:10.1038/hdy.2011.120.

9. Hansen, M.M. Expression of Interest: Transcriptomics and the Designation of Conservation Units. Mol. Ecol. 2010, 19, 1757–1759, doi:10.1111/j.1365-294X.2010.04597.x.

10. Larsen, P.F.; Nielsen, E.E.; Williams, T.D.; Hemmer-Hansen, J.; Chipman, J.K.; Kruhøffer, M.; Grønkjær, P.; George, S.G.; Dyrskjøt, L.; Loeschcke, V. Adaptive Differences in Gene Expression in European Flounder (Platichthys flesus). Mol. Ecol. 2007, 16, 4674–4683, doi:10.1111/j.1365-294X.2007.03530.x.

11. Whitehead, A.; Crawford, D.L. Variation within and among Species in Gene Expression: Raw Material for Evolution. Mol. Ecol. 2006, 15, 1197–1211, doi:10.1111/j.1365-294X.2006.02868.x.

12. Passow, C.N.; Henpita, C.; Shaw, J.H.; Quackenbush, C.R.; Warren, W.C.; Schartl, M.; Arias-Rodriguez, L.; Kelley, J.L.; Tobler, M. The Roles of Plasticity and Evolutionary Change in Shaping Gene Expression Variation in Natural Populations of Extremophile Fish. Mol. Ecol. 2017, 26, 6384–6399, doi:10.1111/mec.14360.

13. Etges, W.J.; de Oliveira, C.C.; Rajpurohit, S.; Gibbs, A.G. Effects of Temperature on Transcriptome and Cuticular Hydrocarbon Expression in Ecologically Differentiated Populations of Desert Drosophila. Ecol. Evol. 2017, 7, 619–637, doi:10.1002/ece3.2653.

14. He, X.; Wilson, C.C.; Wellband, K.W.; Houde, A.L.S.; Neff, B.D.; Heath, D.D. Transcriptional Profiling of Two Atlantic Salmon Strains: Implications for Reintroduction into Lake Ontario. Conserv. Genet. 2015, 16, 277– 287, doi:10.1007/s10592-014-0657-1.

15. Tymchuk, W.; O’Reilly, P.; Bittman, J.; MacDonald, D.; Schulte, P. Conservation Genomics of Atlantic Salmon: Variation in Gene Expression between and within Regions of the Bay of Fundy. Mol. Ecol. 2010, 19, 1842– 1859, doi:10.1111/j.1365-294X.2010.04596.x.

16. Quinn, N.L.; Mcgowan, C.R.; Cooper, G.A.; Koop, B.F.; Davidson, W.S. Identification of Genes Associated with Heat Tolerance in Arctic Charr Exposed to Acute Thermal Stress. Physiol Genomics 2011, 43, 685–696, doi:10.1152/physiolgenomics.00008.2011.-Arctic.

17. Allen, S.L.; Bonduriansky, R.; Sgro, C.M.; Chenoweth, S.F. Sex-Biased Transcriptome Divergence along a Latitudinal Gradient. Mol. Ecol. 2017, 26, 1256–1272, doi:10.1111/mec.14015.

18. De Frenne, P.; Graae, B.J.; Rodríguez-Sánchez, F.; Kolb, A.; Chabrerie, O.; Decocq, G.; De Kort, H.; De Schrijver, A.; Diekmann, M.; Eriksson, O.; et al. Latitudinal Gradients as Natural Laboratories to Infer Species’ Responses to Temperature. J. Ecol. 2013, 101, 784–795, doi:10.1111/1365-2745.12074.

19. Juneja, P.; Quinn, A.; Jiggins, F.M. Latitudinal Clines in Gene Expression and Cis-Regulatory Element Variation in Drosophila melanogaster. BMC Genomics 2016, 17, 1–11, doi:10.1186/s12864-016-3333-7.

20. Pörtner, H.O.; Bock, C.; Knust, R.; Lannig, G.; Lucassen, M.; Mark, F.C.; Sartoris, F.J. Cod and Climate in a Latitudinal Cline: Physiological Analyses of Climate Effects in Marine Fishes. Clim. Res. 2008, 37, 253–270, doi:10.3354/cr00766.

21. Schmidt, P.S.; Serrão, E.A.; Pearson, G.A.; Riginos, C.; Rawson, P.D.; Hilbish, T.J.; Brawley, S.H.; Trussell, G.C.; Carrington, E.; Wethey, D.S.; et al. Ecological Genetics in the North Atlantic: Environmental Gradients and Adaptation at Specific Loci. Ecology 2008, 89, 91–107, doi:10.1890/07-1162.1.

22. Avise, J.C. Phylogeography: Retrospect and Prospect. J. Biogeogr. 2009, 36, 3–15, doi:10.1111/j.1365-2699.2008.02032.x.

23. Moritz, C.; Langham, G.; Kearney, M.; Krockenberger, A.; vanDerWal, J.; Williams, S. Integrating Phylogeography and Physiology Reveals Divergence of Thermal Traits between Central and Peripheral Lineages of Tropical Rainforest Lizards. Philos. Trans. R. Soc. B Biol. Sci. 2012, 367, 1680–1687, doi:10.1098/rstb.2012.0018.

24. Jueterbock, A.; Franssen, S.U.; Bergmann, N.; Gu, J.; Coyer, J.A.; Reusch, T.B.H.; Bornberg-Bauer, E.; Olsen, J.L. Phylogeographic Differentiation versus Transcriptomic Adaptation to Warm Temperatures in Zostera marina, a Globally Important Seagrass. Mol. Ecol. 2016, In Press, 5396–5411, doi:10.1111/mec.13829.

25. da Costa, F.; Cerviño-Otero, A.; Iglesias, Ó.; Cruz, A.; Guévélou, E. Hatchery Culture of European Clam Species (Family Veneridae). Aquac. Int. 2020, 28, 1675–1708, doi:10.1007/s10499-020-00552-x.

26. Borsa, P.; Millet, B. Recruitment of the Clam Ruditapes decussatus in the Lagoon of Thau, Mediterranean. Estuar. Coast. Shelf Sci. 1992, 35, 289–300, doi:10.1016/S0272-7714(05)80049-6.

27. Gharbi, A.; Chatti, N.; Said, K.; Wormhoudt, A. Genetic Variation and Population Structure of the Carpet Shell Clam Ruditapes decussatus along the Tunisian Coast Inferred from MtDNA and ITS1 Sequence Analysis. Biologia (Bratisl). 2010, 65, 688–696, doi:10.2478/s11756-010-0069-8.

28. Habtemariam, B.T.; Arias, A.; García-Vázquez, E.; Borrell, Y.J. Impacts of Supplementation Aquaculture on the Genetic Diversity of Wild Ruditapes decussatus from Northern Spain. Aquac. Environ. Interact. 2015, 6, 241–254, doi:10.3354/aei00128.

29. Leite, R.B.; Afonso, R.; Cancela, M.L. Perkinsus Sp. Infestation in Carpet-Shell Clams, Ruditapes decussatus (L), along the Portuguese Coast. Results from a 2-Year Survey. Aquaculture 2004, 240, 39–53, doi:10.1016/j.aquaculture.2004.05.014.

30. Cordero, D.; Peña, J.B.; Saavedra, C. Phylogeographic Analysis of Introns and Mitochondrial DNA in the Clam Ruditapes decussatus Uncovers the Effects of Pleistocene Glaciations and Endogenous Barriers to Gene Flow. Mol. Phylogenet. Evol. 2014, 71, 274–287, doi:10.1016/j.ympev.2013.11.003.

31. Arias-Pérez, A.; Cordero, D.; Borrell, Y.; Sánchez, J.A.; Blanco, G.; Freire, R.; Insua, A.; Saavedra, C. Assessing the Geographic Scale of Genetic Population Management with Microsatellites and Introns in the Clam Ruditapes decussatus. Ecol. Evol. 2016, 6, doi:10.1002/ece3.2052.

32. Markaide, P.; Gairín, I.; Cordero, D.; Ibarrola, I.; Saavedra, C. No Hybridization and Marked Interspecific Differences in Individual Growth Rate in Mixed Cultures of Manila Clam (Ruditapes philippinarum) and Grooved Carpet-Shell Clam (R. decussatus). Aquaculture 2021, 541, 736824, doi:10.1016/j.aquaculture.2021.736824.

33. Pascual, M.; Rives, B.; Schunter, C.; MaCpherson, E. Impact of Life History Traits on Gene Flow: A Multispecies Systematic Review across Oceanographic Barriers in the Mediterranean Sea. PLoS One 2017, 12, doi:10.1371/JOURNAL.PONE.0176419.

34. Patarnello, T.; Volckaert, F.A.M.J.; Castilho, R. Pillars of Hercules: Is the Atlantic-Mediterranean Transition a Phylogeographical Break? Mol. Ecol. 2007, 16, 4426–4444, doi:10.1111/j.1365-294X.2007.03477.x.

35. Rousset, F. GENEPOP’007: A Complete Re-Implementation of the GENEPOP Software for Windows and Linux. Mol. Ecol. Resour. 2008, 8, 103–106, doi:10.1111/j.1471-8286.2007.01931.x.

36. Falush, D.; Stephens, M.; Pritchard, J. Inference of Population Structure Using Multilocus Genotype Data: Linked Loci and Correlated Allele Frequencies. Genetics 2003, 164, 1567–1587, doi:10.1111/j.1471-8286.2007.01758.x.

37. n, S.; Nei, M. The Neighbor-Joining Method: A New Method for Reconstructing Phylogenetic Trees. Mol. Biol. Evol. 1987, 4, 406–425, doi:citeulike-article-id:93683.

38. Leite, R.B.; Milan, M.; Coppe, A.; Bortoluzzi, S.; dos Anjos, A.; Reinhardt, R.; Saavedra, C.; Patarnello, T.; Cancela, M.L.; Bargelloni, L. MRNA-Seq and Microarray Development for the Grooved Carpet Shell Clam, Ruditapes decussatus: A Functional Approach to Unravel Host -Parasite Interaction. BMC Genomics 2013, 14, doi:10.1186/1471-2164-14-741.

39. Milan, M.; Coppe, A.; Reinhardt, R.; Cancela, L.M.; Leite, R.B.; Saavedra, C.; Ciofi, C.; Chelazzi, G.; Patarnello, T.; Bortoluzzi, S.; et al. Transcriptome Sequencing and Microarray Development for the Manila Clam, Ruditapes philippinarum: Genomic Tools for Environmental Monitoring. BMC Genomics 2011, 12, 234, doi:10.1186/1471-2164-12-234.

40. Saeed, A.I.; Sharov, V.; White, J.; Li, J.; Liang, W.; Bhagabati, N.; Braisted, J.; Klapa, M.; Currier, T.; Thiagarajan, M.; et al. TM4: A Free, Open-Source System for Microarray Data Management and Analysis. Biotechniques 2003, 34, 374–378, doi:10.2144/03342MT01.

41. Benjamini, Y.; Hochberg, Y. Controlling the False Discovery Rate: A Practical and Powerful Approach to Multiple Testing. J. R. Stat. Soc. Ser. B 1995, 57, 289–300, doi:10.1111/J.2517-6161.1995.TB02031.X.

42. Eisen, M.B.; Spellman, P.T.; Brown, P.O.; Botstein, D. Cluster Analysis and Display of Genome-Wide Expression Patterns. Genetics 1998, 95, 14863–14868.

43. Conesa, A.; Götz, S.; García-Gómez, J.M.; Terol, J.; Talón, M.; Robles, M. Blast2GO: A Universal Tool for Annotation, Visualization and Analysis in Functional Genomics Research. Bioinformatics 2005, 21, 3674–3676, doi:10.1093/bioinformatics/bti610.

44. Huang, D.W.; Lempicki, R. a; Sherman, B.T. Systematic and Integrative Analysis of Large Gene Lists Using DAVID Bioinformatics Resources. Nat. Protoc. 2009, 4, 44–57, doi:10.1038/nprot.2008.211.

45. Rohlfs, R. V.; Nielsen, R. Phylogenetic ANOVA: The Expression Variance and Evolution Model for Quantitative Trait Evolution. Syst. Biol. 2015, 64, 695–708, doi:10.1093/SYSBIO/SYV042.

46. Gleason, L.U.; Burton, R.S. RNA-Seq Reveals Regional Differences in Transcriptome Response to Heat Stress in the Marine Snail Chlorostoma funebralis. Mol. Ecol. 2015, 24, 610–627, doi:10.1111/mec.13047.

47. Hughes, D.A.; Kircher, M.; He, Z.; Guo, S.; Fairbrother, G.L.; Moreno, C.S.; Khaitovich, P.; Stoneking, M. Evaluating Intra- and Inter-Individual Variation in the Human Placental Transcriptome. Genome Biol. 2015, 16, 54, doi:10.1186/s13059-015-0627-z.

48. Whitehead, A.; Crawford, D.L. Neutral and Adaptive Variation in Gene Expression. Proc. Natl. Acad. Sci. U. S. A. 2006, 103, 5425–5430, doi:10.1073/pnas.0507648103.

49. Thiagalingam, S. Epigenetic Memory in Development and Disease: Unraveling the Mechanism. Biochim. Biophys. Acta - Rev. Cancer 2020, 1873.

50. Wang, X.; Li, A.; Wang, W.; Que, H.; Zhang, G.; Li, L. DNA Methylation Mediates Differentiation in Thermal Responses of Pacific Oyster (Crassostrea gigas) Derived from Different Tidal Levels. Heredity (Edinb). 2020, 1–13, doi:10.1038/s41437-020-0351-7.

51. Zhang, X.; Li, Q.; Kong, L.; Yu, H. Epigenetic Variation of Wild Populations of the Pacific Oyster Crassostrea gigas Determined by Methylation-Sensitive Amplified Polymorphism Analysis. Fish. Sci. 2018, 84, 61– 70, doi:10.1007/s12562-017-1154-5.

52. Schunter, C.; Carreras-Carbonell, J.; MacPherson, E.; TintorÉ, J.; Vidal-Vijande, E.; Pascual, A.; Guidetti, P.; Pascual, M. Matching Genetics with Oceanography: Directional Gene Flow in a Mediterranean Fish Species. Mol. Ecol. 2011, 20, 5167–5181, doi:10.1111/j.1365-294X.2011.05355.x.

53. Lahtvee, P.-J.; Sá Nchez, B.J.; Smialowska, A.; Kasvandik, S.; Elsemman, I.E.; Gatto, F.; Nielsen, J. Absolute Quantification of Protein and MRNA Abundances Demonstrate Variability in Gene-Specific Translation Efficiency in Yeast. Cell Syst. 2017, 4, 495–504.e5, doi:10.1016/j.cels.2017.03.003.

54. Yon Rhee, S.; Wood, V.; Dolinski, K.; Draghici, S. Use and Misuse of the Gene Ontology Annotations. Nat. Rev. Genet. 2008 97 2008, 9, 509–515, doi:10.1038/nrg2363.

55. Petibon, C.; Malik Ghulam, M.; Catala, M.; Abou Elela, S. Regulation of Ribosomal Protein Genes: An Ordered Anarchy. Wiley Interdiscip. Rev. RNA 2021, 12, 1–29, doi:10.1002/wrna.1632.

56. Blanchette, B.; Feng, X.; Singh, B.R. Marine Glutathione S-Transferases. Mar. Biotechnol. 2007, 9, 513–542, doi:10.1007/s10126-007-9034-0.

57. Hayes, J.D.; Flanagan, J.U.; Jowsey, I.R. Glutathione transferases. Annu. Rev. Pharmacol. Toxicol. 2005, 45, 51–88, doi:10.1146/annurev.pharmtox.45.120403.095857.

58. Hellou, J.; Ross, N.W.; Moon, T.W. Glutathione, Glutathione S-Transferase, and Glutathione Conjugates, Complementary Markers of Oxidative Stress in Aquatic Biota. Environ. Sci. Pollut. Res. 2012, 19, 2007–2023, doi:10.1007/s11356-012-0909-x.

59. Verlecar, X.N.; Jena, K.B.; Chainy, G.B.N. Seasonal Variation of Oxidative Biomarkers in Gills and Digestive Gland of Green-Lipped Mussel Perna viridis from Arabian Sea. Estuar. Coast. Shelf Sci. 2008, 76, 745– 752, doi:10.1016/j.ecss.2007.08.002.

60. Kim, M.; Ahn, I.Y.; Cheon, J.; Park, H. Molecular Cloning and Thermal Stress-Induced Expression of a Pi-Class Glutathione S-Transferase (GST) in the Antarctic Bivalve Laternula elliptica. Comp. Biochem. Physiol. - A Mol. Integr. Physiol. 2009, 152, 207–213, doi:10.1016/j.cbpa.2008.09.028.

61. Anacleto, P.; Maulvault, A.L.; Lopes, V.M.; Repolho, T.; Diniz, M.; Nunes, M.L.; Marques, A.; Rosa, R. Ecophysiology of Native and Alien-Invasive Clams in an Ocean Warming Context. Comp. Biochem. Physiol. -Part A Mol. Integr. Physiol. 2014, 175, 28–37, doi:10.1016/j.cbpa.2014.05.003.

62. Junge, W.; Nelson, N. ATP Synthase. Annu. Rev. Biochem. 2015, 84, 631–657, doi:10.1146/annurev-biochem-060614-034124.

63. Itoi, S.; Kinoshita, S.; Kikuchi, K.; Watabe, S. Changes of Carp FoF1-ATPase in Association with Temperature Acclimation. Am. J. Physiol. - Regul. Integr. Comp. Physiol. 2003, 284, 153–163, doi:10.1152/ajpregu.00182.2002.

64. Kikuchi, K.; Itoi, S.; Watabe, S. Increased Levels of Mitochondrial ATP Synthase β-Subunit in Fast Skeletal Muscle of Carp Acclimated to Cold Temperature. Fish. Sci. 1999, 65, 629–636, doi:10.2331/FISHSCI.65.629.

65. Chenghua Li Isolation of an ATP Synthase CDNA from Sinonovacula constricta and Its MRNA Expression by Thermal Stress. African J. Biotechnol. 2012, 11, 1719–1723, doi:10.5897/ajb11.076.

66. Fields, P.A.; Eraso, A. A Year in the Salt Marsh: Seasonal Changes in Gill Protein Expression in the Temperate Intertidal Mussel Geukensia demissa. Mar. Environ. Res. 2020, 161, 105088, doi:10.1016/j.marenvres.2020.105088.

67. Wijers, S.L.J.; Smit, E.; Saris, W.H.M.; Mariman, E.C.M.; Van Marken Lichtenbelt, W.D. Cold- and Overfeeding-Induced Changes in the Human Skeletal Muscle Proteome. J. Proteome Res. 2010, 9, 2226–2235, doi:10.1021/pr9010074.

68. Coppe, A.; Agostini, C.; Marino, I.A.M.; Zane, L.; Bargelloni, L.; Bortoluzzi, S.; Patarnello, T. Genome Evolution in the Cold: Antarctic Icefish Muscle Transcriptome Reveals Selective Duplications Increasing Mitochondrial Function. Genome Biol. Evol. 2013, 5, 45–60, doi:10.1093/gbe/evs108.

69. Price, P.D.; Palmer Droguett, D.H.; Taylor, J.A.; Kim, D.W.; Place, E.S.; Rogers, T.F.; Mank, J.E.; Cooney, C.R.; Wright, A.E. Detecting Signatures of Selection on Gene Expression. Nat. Ecol. Evol. 2022, 6, 1035– 1045, doi:10.1038/s41559-022-01761-8.

70. Sussarellu, R.; Huvet, A.; Lapègue, S.; Quillen, V.; Lelong, C.; Cornette, F.; Jensen, L.F.; Bierne, N.; Boudry, P. Additive Transcriptomic Variation Associated with Reproductive Traits Suggest Local Adaptation in a Recently Settled Population of the Pacific Oyster, Crassostrea gigas. BMC Genomics 2015, 16, 808, doi:10.1186/s12864-015-1972-8.

71. Brauer, C.J.; Unmack, P.J.; Beheregaray, L.B. Comparative Ecological Transcriptomics and the Contribution of Gene Expression to the Evolutionary Potential of a Threatened Fish. Mol. Ecol. 2017, 26, 6841– 6856, doi:10.1111/MEC.14432.

