## Supplementary Table and Figures for "Transcriptional profiling of populations in the clam *Ruditapes decussatus* suggests genetically determined differentiation in gene-expression along parallel temperature gradients and between races of the Atlantic Ocean and West Mediterranean sea"

Table

Table S1. Allelic frequencies in the four sampled populations for the six intron RFLP markers scored.

Figures

Figure S1. Results of Bayesian analysis of genetic structure based on allelic frequencies of six intron RFLP genetic markers for K=3

Figure S2. Principal components analysis of expression data from all samples

Figure S3. Heatmap showing gene expression variation across samples for the 105 genes which were significant for the Race factor in the 2-way ANOVA and contributed to the associated enriched GO terms.

Figure S4. Heatmap showing gene expression variation across samples for the genes which were significant for Latitude only in the 2-way ANOVA and contributed to the enriched GO terms.

Figure S5. Average expression in populations for the 9 genes which were significant for Race x Latitude interaction in the 2-way ANOVA

Table S1. Allelic frequencies in the four sampled populations for the six intron RFLP markers scored. N is the sample size.

| Locus | Allele | Locality | | | |
|---|---|---|---|---|---|
| | | AN | AS | MS | MN |
| CL116-ia | A | 0.667 | 0.679 | 0.789 | 0.732 |
| | B | 0.333 | 0.321 | 0.211 | 0.268 |
| | N | 24 | 56 | 19 | 41 |
| CL13-ia | A | 0.958 | 0.797 | 0.842 | 0.922 |
| | B | 0.042 | 0.203 | 0.158 | 0.078 |
| | N | 24 | 59 | 19 | 45 |
| CL102-ia | A | 0.292 | 0.682 | 0.611 | 0.257 |
| | B | 0.708 | 0.318 | 0.389 | 0.743 |
| | N | 24 | 55 | 18 | 35 |
| CL126-ia | A | 0.667 | 0.745 | 0.289 | 0.128 |
| | B | 0.333 | 0.255 | 0.605 | 0.802 |
| | C | 0.000 | 0.000 | 0.105 | 0.058 |
| | D | 0.000 | 0.000 | 0.000 | 0.012 |
| | N | 21 | 51 | 19 | 43 |
| TBP | A | 0.283 | 0.202 | 0.553 | 0.677 |
| | B | 0.717 | 0.798 | 0.447 | 0.290 |
| | C | 0.000 | 0.000 | 0.000 | 0.032 |
| | D | - | - | - | - |
| | N | 23 | 57 | 19 | 31 |
| SRP54 | A | 0.250 | 0.114 | 0.368 | 0.361 |
| | B | 0.568 | 0.737 | 0.632 | 0.583 |
| | C | 0.091 | 0.053 | 0.000 | 0.028 |
| | D | 0.091 | 0.096 | 0.000 | 0.028 |
| | N | 22 | 57 | 19 | 36 |

Figure S1. Results of Bayesian analysis of genetic structure based on allelic frequencies of six intron RFLP genetic markers (Cordero et al, 2014). Data from the four sampled localities for this study (marked above the chart) were analyzed together with data from 11 other localities from a previous study by Cordero et al (2014). The three races described by Cordero et al (2014) are indicated below the chart with colored lines.

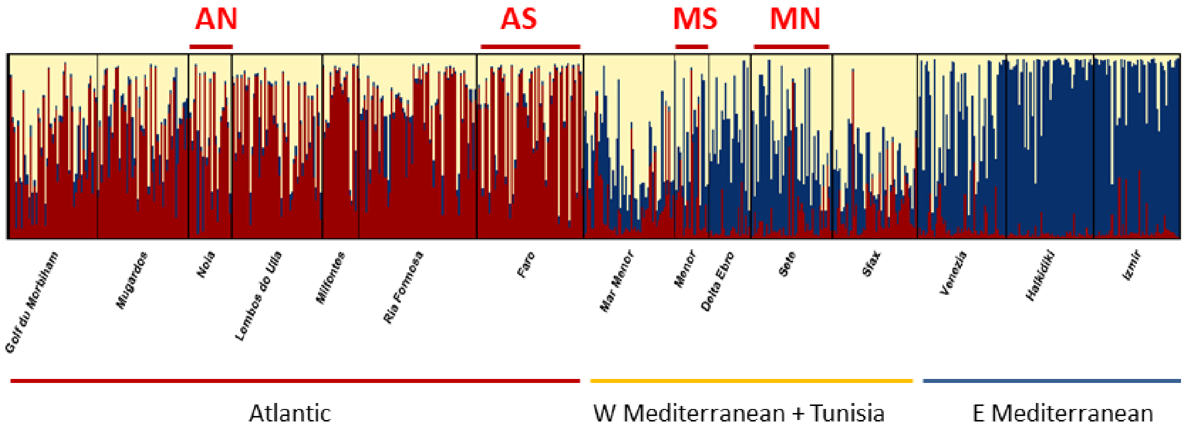

Figure S2. Principal components analysis of expression data from all samples.

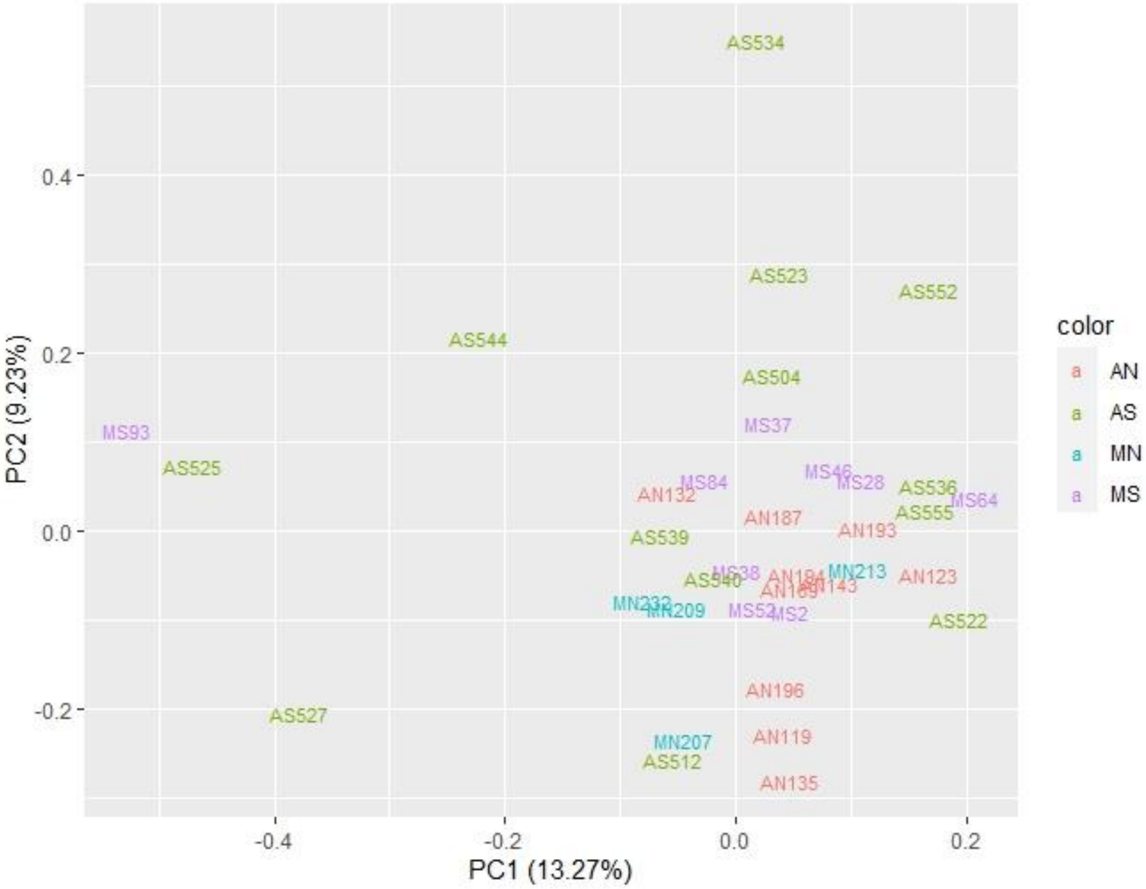

Figure S3. Heatmap showing gene expression variation across samples for the 105 genes which were significant for the Race factor in the 2-way ANOVA and contributed to the associated enriched GO terms.

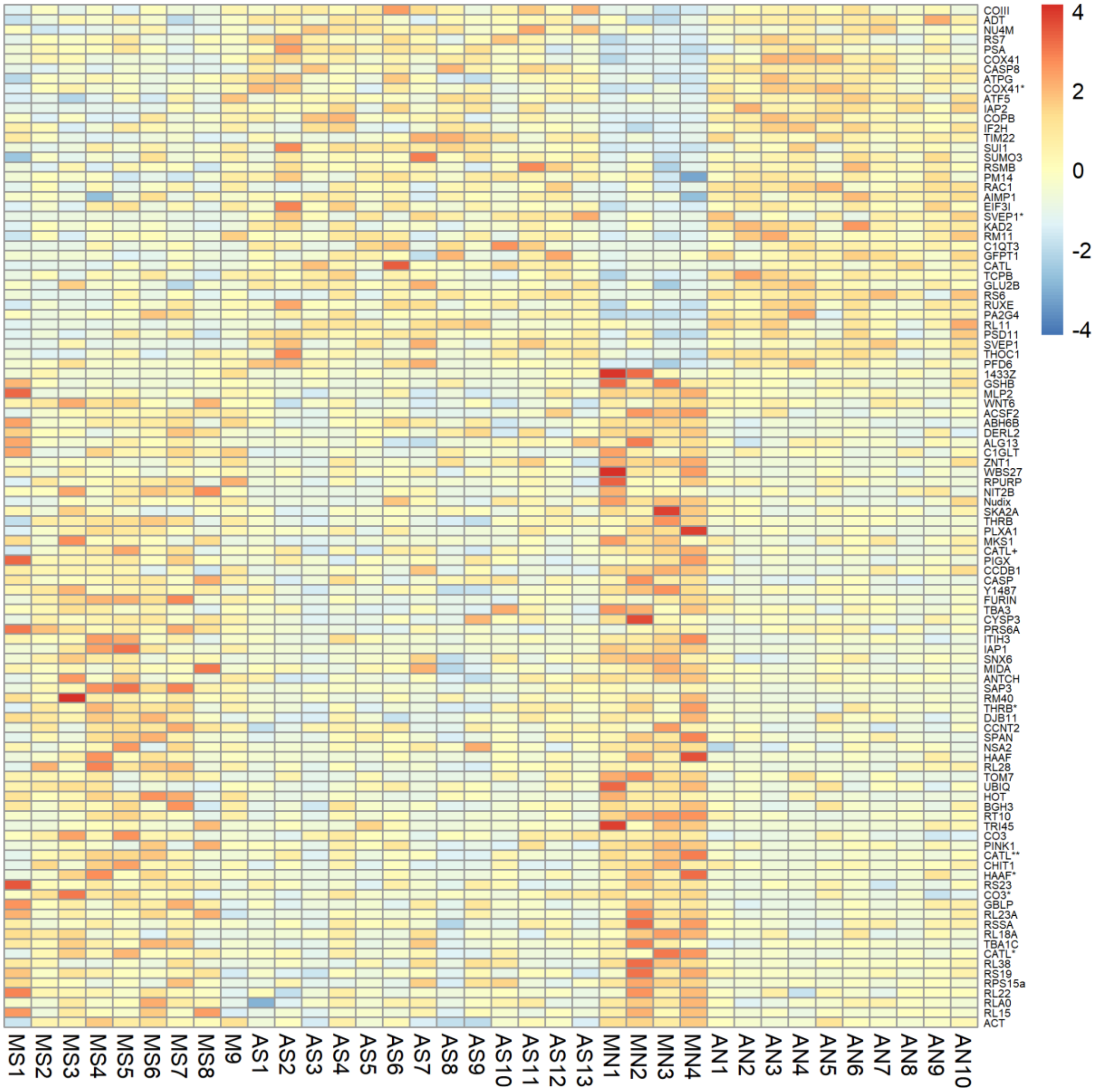

Figure S4. Heatmap showing gene expression variation across samples for the genes that were significant for Latitude only in the 2-way ANOVA and contributed to the enriched GO terms.

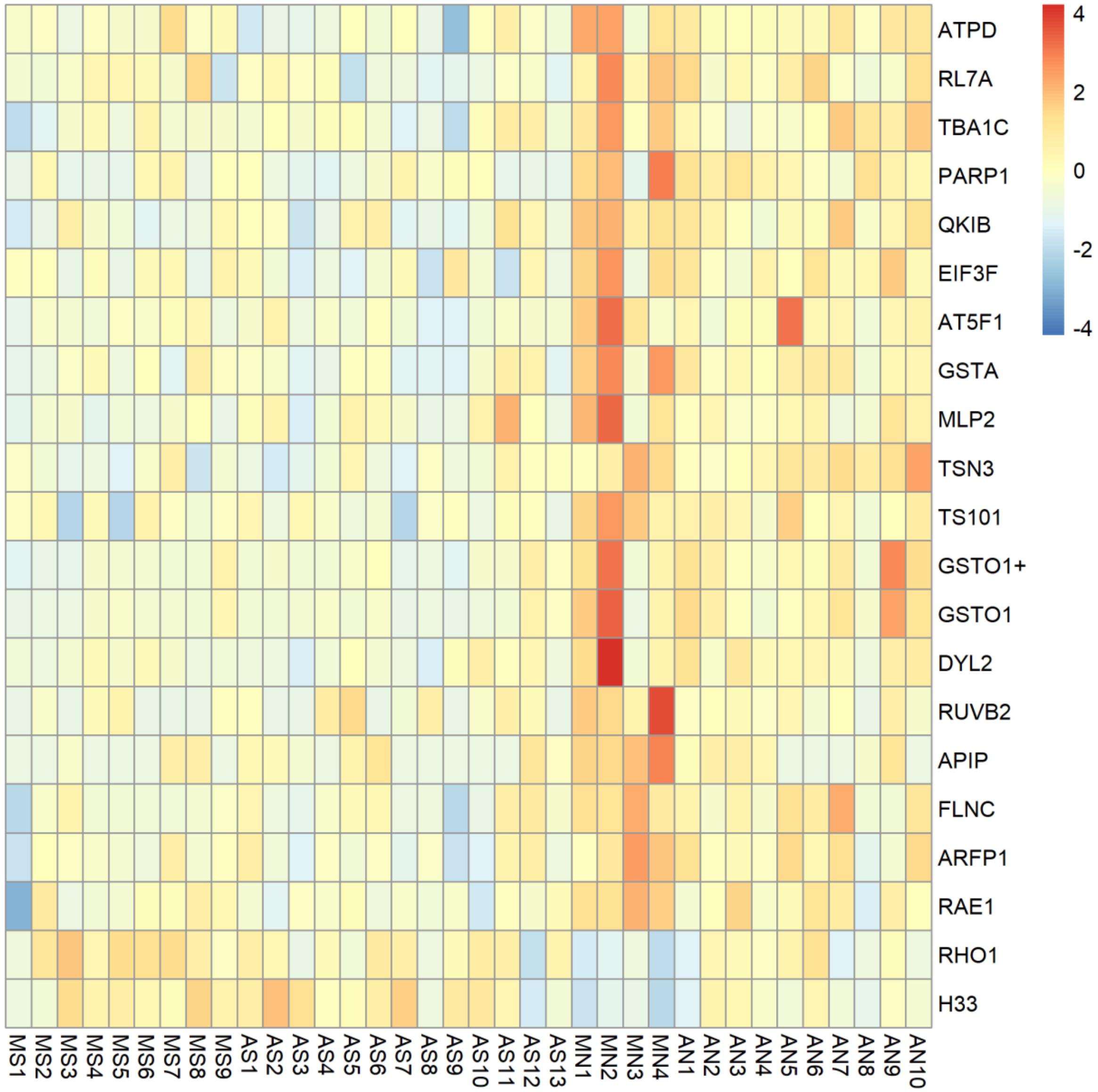

FigureS5.- Average expression in populations for the 9 genes that were significant for Race x Latitude interaction in the 2-way ANOVA. Note that the normalized signal intensity is in logarithmic scale. Gene numbers (#) as in supplementary file 1.

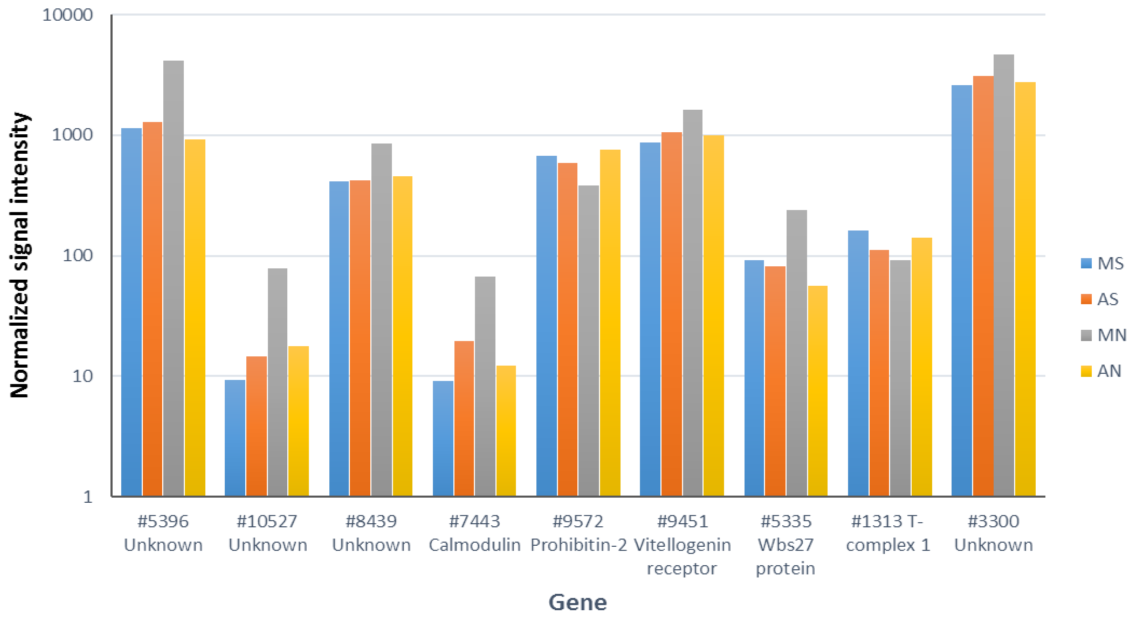
